# High-fidelity dendritic sodium spike generation in human layer 2/3 neocortical pyramidal neurons

**DOI:** 10.1101/2022.03.16.484669

**Authors:** Helen M. Gooch, Tobias Bluett, Madhusoothanan B. Perumal, Hong D. Vo, Lee N. Fletcher, Jason Papacostas, Rosalind L. Jeffree, Martin Wood, Michael J. Colditz, Jason McMillen, Tony Tsahtsarlis, Damian Amato, Robert Campbell, Lisa Gillinder, Stephen R. Williams

## Abstract

Dendritic computations have a central role in rodent neocortical circuit function, which are signaled into neuronal output by the initiation and propagation of regenerative dendritic spikes. However, it remains poorly explored whether these mechanisms are operational in the human neocortex. To directly examine the dendritic computational properties of the most numerous class of pyramidal neuron in the human and rat neocortex we made simultaneous electrical recordings from the soma and apical dendrites of layer 2/3 pyramidal neurons maintained in acute brain slices of analogous cortical areas under identical experimental conditions. In both species correlated dendritic excitatory input led to the initiation of sodium channel-mediated dendritic spikes with conserved biophysical properties. Dendritic sodium spikes in human and rat layer 2/3 pyramidal neurons could be generated across a similar wide input range, exhibited a similar frequency range of activation, and forward propagated with high fidelity to drive neuronal output to implement stereotyped computations. Our findings therefore reveal that the dendritic computational properties of human layer 2/3 pyramidal neurons are phylogenetically conserved.

## Introduction

The human neocortical mantle is a highly evolved structure ^1^, made up of hundreds of functionally and anatomically defined areas ^2^. Although the neuronal classes which form local neocortical circuits are largely preserved across mammalian species ^3,4^, the dendritic tree of excitatory pyramidal neurons of the human neocortex are morphologically elaborated and harbour a greater number of dendritic spines than those of homologous neurons in rodents and other primates ^3,5,6^. In the rodent neocortex, the dendrites of pyramidal neurons are electrically active and powerfully influence neuronal output by the generation of regenerative dendritic spikes ^7–11^, which are engaged during *in vivo* circuit computations ^12–21^, to enhance single neuron computational capacity. Elaboration of the dendritic tree of human neocortical pyramidal neurons may therefore be one of the factors underlying the enriched computational capacity of the human neocortex.

Pioneering dendritic recordings of human layer 2/3 (HL2/3) and layer 5 (HL5) neocortical pyramidal neurons have revealed that human dendrites are electrically excitable and capable of initiating regenerative dendritic spikes ^22–24^. Surprisingly, however, the properties of dendritic electrical excitability and the computational operations executed in the dendritic trees of human neocortical pyramidal neurons have been reported to diverge from those of rodents ^22–24^. Specifically, simultaneous recordings from the apical dendrites of human HL5 pyramidal neurons have shown that dendritic spikes are highly compartmentalized within the apical dendritic tree, and have a minimal impact on action potential (AP) generation ^24^, in contrast to the well-characterized powerful forward propagation of apical dendritic spikes in rodent L5B pyramidal neurons that drive AP burst firing ^7,8,11,24–26^ Recent work has, however, suggested that this disparity may be influenced by the experimental sampling of distinct classes of L5 pyramidal neurons between species, with significant dendritic calcium electrogenesis found in molecularly identified subcortically projecting L5 pyramidal neurons of both taxa ^27^.

Notably, it has recently been reported that the dendritic integrative operations of HL2/3 pyramidal neurons are mechanistically and functionally divergent from those of the rodent, with direct recordings demonstrating the initiation and forward propagation of calcium channel-mediated apical dendritic spikes ^22^, a dendritic spike generation mechanism not apparent in rodent L2/3 pyramidal neurons ^10,14,28,29^. This fundamental species-specific difference in active dendritic integration has been suggested to transform the dendritic computations implemented by HL2/3 pyramidal neurons, allowing the execution of anti-coincidence computations, as calcium channel-mediated dendritic spikes were found to be activated only within a narrow range of dendritic excitatory input ^22^. As such anti-coincidence computations are distinct from the correlative computations executed throughout the dendritic trees of rodent pyramidal neurons ^8,28,30–33^ that have prominent roles in neuronal circuit operations in the rodent neocortex ^12–15,17,18,21,34^, and other circuits ^35–37^, we sought to compare the dendritic integrative properties of L2/3 pyramidal neurons of the human temporal cortex with those of an analogous neocortical area of the rodent under identical experimental conditions. We find that dendritic excitatory input in human and rat L2/3 pyramidal neurons is locally integrated at dendritic sites, but surprisingly dendritic spikes in both species are mediated by dendritic sodium channels. Dendritic sodium spikes could be generated across a wide range of correlated excitatory input, at high repetition frequencies, and powerfully forward propagated to drive neuronal output in both human and rat L2/3 pyramidal neurons to implement stereotyped computations. Thus, our findings reveal that the computational capacity of HL2/3 pyramidal neurons is not transformed by a divergence of integrative mechanisms, but is enhanced by the implementation of a phylogenetically conserved dendritic spiking mechanism in the larger and more complex dendritic trees of HL2/3 pyramidal neurons.

## Results

### Morphological and electrical architecture of human and rat L2/3 pyramidal neurons

To examine the electrophysiological and morphological properties of pyramidal neurons of the human and rodent neocortex we prepared acute brain slices of the human temporal cortex, obtained from patients who had undergone neurosurgery for the alleviation of refractory epilepsy, or the removal of sub-cortical brain tumours (Supplementary Tables 1 and 2), and an analogous neocortical area of the rat, the temporal associative neocortex ^24^. High-resolution reconstruction of the dendritic morphology of biocytin-filled pyramidal neurons distributed throughout layers 2 and 3 of the human and rodent temporal cortex revealed that the apical and basal dendritic trees of HL2/3 pyramidal neurons were significantly larger and more complex than those of rat L2/3 (RL2/3) pyramidal neurons (distance from neocortical surface: HL2/3= 824 ± 45 μm, range= 449 - 1203 μm, n= 20 reconstructed neurons; RL2/3= 343 ± 19.5 μm, range= 212 - 540 μm, n= 20 reconstructed neurons; Fig. 1a,b, Supplementary Fig. 1). Quantitative analysis showed that the number of primary dendrites, the path lengths, and the number of branch points in the basal and apical dendritic trees of HL2/3 pyramidal neurons were significantly greater, longer, and more complex than RL2/3 pyramidal neurons (primary dendrites: HL2/3= 7.4 ± 0.3, RL2/3= 5.9 ± 0.3, Mann Whitney U= 81.5, *P*= 0.0006; apical path length= HL2/3= 780.4 ± 14.8 μm, RL2/3= 358.1 ± 7.9 μm, t-test, df= 467, *P*< 0.0001; basal path length: HL2/3= 211.0 ± 3.1 μm, RL2/3= 133.6 ± 2.2 μm, Mann Whitney U= 123035, *P*< 0.0001; apical branch points: HL2/3= 39.7 ± 2.9, RL2/3= 17.3 ± 1.1, Mann Whitney U= 7.5, *P*< 0.0001; basal branch points: HL2/3= 36.8 ± 2.4, RL2/3= 24.2 ± 1.3, t-test, df= 38, *P*< 0.0001; Fig. 1c-h). Furthermore, the calibre of basal and apical dendrites as they emerged from the somata of HL2/3 pyramidal neurons were greater in diameter than in RL2/3 pyramidal neurons; however, the calibre of the main apical dendrite of HL2/3 pyramidal neurons tapered and was similar in diameter to that of RL2/3 pyramidal neurons at the first major bifurcation (Supplementary Fig. 1).

**Figure 1.**
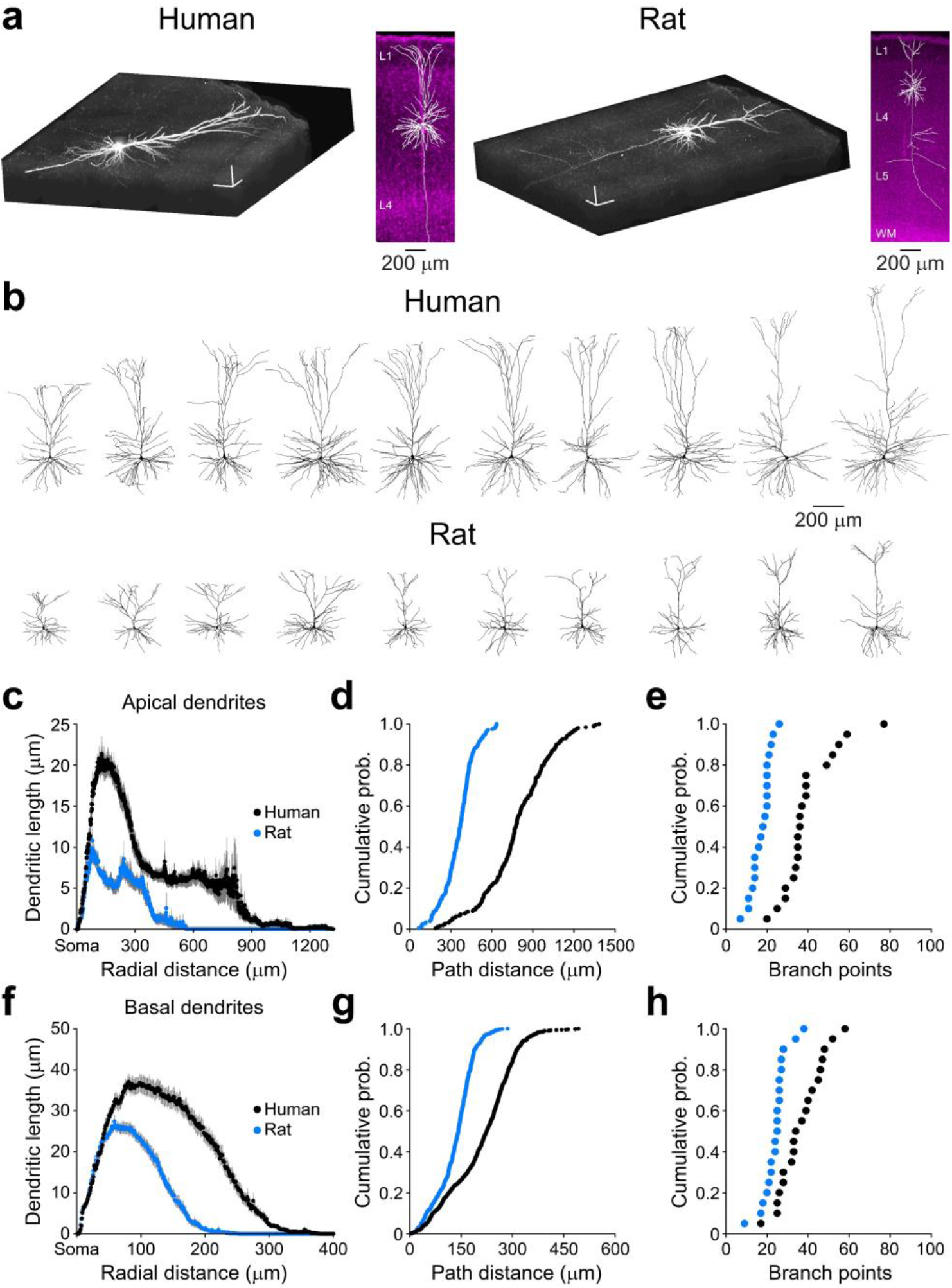
Morphological elaboration of the dendritic tree of human L2/3 pyramidal neurons. **a,** Volume representation of a biocytin-filled HL2/3 and RL2/3 pyramidal neuron. The scale bars represent 100 μm. The insets show reconstructions of the neurons (white) superposed onto low power fluorescence images of neocortical slices counterstained with the nuclear marker DAPI (magenta). The descending axons have been partly reconstructed. **b,** 2-dimensional projections of ten representative reconstructed human and rat L2/3 pyramidal neurons. **c,** Sholl analysis of the apical dendritic tree of HL2/3 (black symbols) and RL2/3 (blue symbols) pyramidal neurons (n= 20 neurons in each species, grey lines represent ± S.E.M.). **d-e,** Cumulative probability distributions of path distance from soma to termination of apical dendrites (d) and the number of branch points in the apical dendritic tree (e). **f,** Sholl analysis of the basal dendritic tree of HL2/3 and RL2/3 pyramidal neurons. **g-h,** Cumulative probability distributions of path distance from soma to termination of basal dendrites (g) and the number of branch points in the basal dendritic tree (h).

How do such differences in the size and complexity of the dendritic tree influence the electrical structure of pyramidal neurons of each species? To experimentally test this, we made simultaneous somatic and apical dendritic whole-cell recordings from human and rat L2/3 pyramidal neurons, and examined the electrical attenuation of simulated excitatory postsynaptic potentials (sEPSPs) as they spread from the somatic or dendritic site of generation through the dendritic tree. Simulated EPSPs were generated by the injection of uniform current waveforms, and exhibited indistinguishable kinetics when generated at and recorded from the soma of human and rat L2/3 pyramidal neurons (driving current: amplitude= 200 pA; τ_rise_= 0.5 ms; τ_decay_= 5 ms; somatic 10-90 % rise-time: human= 3.47 ± 0.10 ms, n= 48, rat= 3.62 ± 0.08 ms, n= 31, t-test, *P*= 0.23; somatic half-width: human= 21.82 ± 0.32 ms, rat= 20.57 ± 0.61 ms, t-test, *P*= 0.08; Fig. 2). To determine the electrical architecture, we first quantified the attenuation of the amplitude of dendritic sEPSPs as they spread from apical dendritic site of generation to the soma (Fig. 2a). In both species the somatic amplitude of dendritically generated sEPSPs progressively decreased as sEPSPs were generated from increasingly remote apical dendritic sites (dendro-somatic voltage transfer, Fig. 2b). To our surprise, when we plotted the degree of dendro-somatic voltage transfer for recordings made from human and rat L2/3 pyramidal neurons against recording distance from the soma, the data fell on a similar curve, which could be fit with an exponential function (length constant= 154 μm, n= 48 HL2/3, n= 31 RL2/3; Fig. 2b). Similarly, when we recorded the somato-dendritic voltage transfer of somatically generated sEPSPs (Fig. 2a,c), or calculated the somatic impact of dendritic sEPSPs (Fig. 2a,d) we found distance-dependent relationship which described similar curves in human and rat L2/3 pyramidal neurons (length constants, 641 μm, and 642 μm, respectively). These data suggest that the attenuation of sEPSPs in the apical dendritic tree of human and rat L2/3 pyramidal neurons is governed by the physical distance between somatic and dendritic recording sites, and does not exhibit species-specific characteristics. Consistent with this interpretation, analysis of dendritic recordings made at comparable physical distances from the soma of human and rat L2/3 pyramidal neurons revealed that the degree of voltage attenuation, and the somatic impact of dendritic sEPSPs, were not statistically different (human distance range= 80 - 192 μm, n= 21; rat distance range= 80 - 191 μm, n= 25; soma-dendrite transfer: human= 0.81 ± 0.02, rat= 0.79 ± 0.02, t-test, *P*= 0.39; dendrite-soma transfer= human= 0.56 ± 0.03, rat= 0.46 ± 0.05, t-test, *P*= 0.07; somatic impact: human= 0.80 ± 0.02, rat= 0.78 ± 0.02, t-test, *P*= 0.31). Furthermore, the pattern of somato-dendritic attenuation of voltage responses generated by the steps of negative current injected at somatic sites in rat and human L2/3 pyramidal neurons described a similar relationship (amplitude= −100pA; length constant= 708 μm; Fig. 2j). To explore if this relationship also held for the electrical filtering of the time-course of synaptic events, we analysed the somatic rise-time of dendritically generated sEPSPs, finding a distance-dependent slowing of the somatic rise-time as sEPSPs were generated from increasingly remote dendritic sites in human and rat L2/3 pyramidal neurons (Supplementary Fig. 2). Analysis revealed that this relationship also had a similar structure between species, and consequently the somatic rise-times of sEPSPs generated from comparable dendritic sites were similar between human and rat L2/3 pyramidal neurons (human distance range= 80 - 192 μm, n= 21; rat distance range= 80 - 191 μm, n= 25, human 10-90 % rise-time= 4.60 ± 0.11, rat 10-90 % rise-time= 4.72 ± 0.09; t-test, *P*= 0.42; Supplementary Fig. 2).

**Figure 2.**
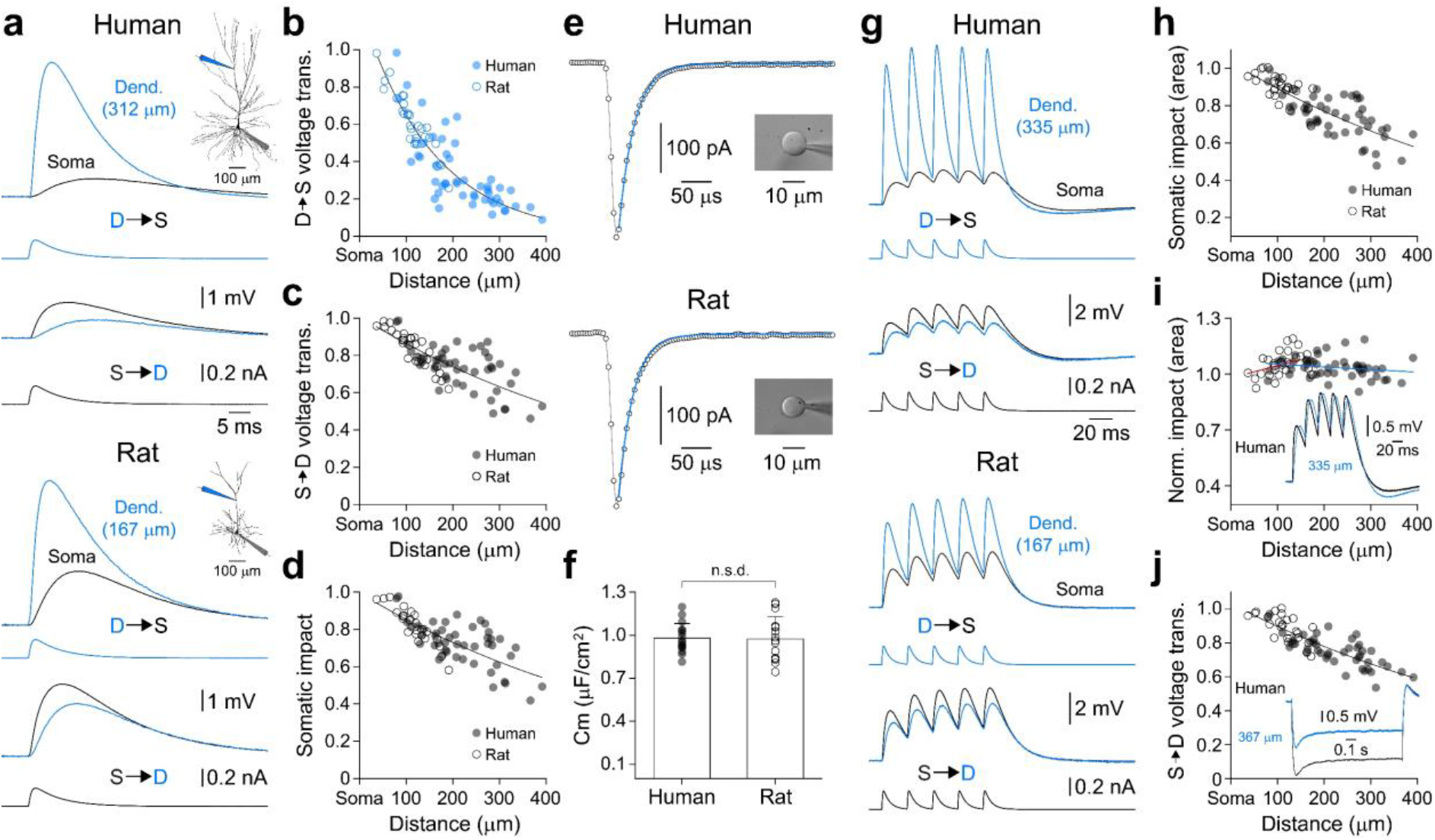
Voltage attenuation in the apical dendritic tree of human and rat L2/3 pyramidal neurons. **a,** Simultaneous dendritic (Dend., blue) and somatic (black) recording of simulated EPSPs (sEPSPs), showing dendro-somatic (D-S) and somato-dendritic (S-D) attenuation in human and rat L2/3 pyramidal neurons. The driving currents injected at dendritic and somatic sites are shown below voltage traces. The morphology of recorded neurons and placement of recording electrodes are inset. **b,** Distance-dependency of sEPSP dendro-somatic voltage transfer in HL2/3 (filled symbols) and RL2/3 (open symbols) pyramidal neurons. The line represents a single exponential fit, with a length constant of 154 μm. **c,** Distance-dependency of sEPSP somato-dendritic voltage transfer in human and rat L2/3 pyramidal neurons. The line represents a single exponential fit with a length constant of 641 μm. **d,** Somatic impact of dendritic sEPSPs, calculated by the division of the somatic amplitude of sEPSPs generated at the soma and apical dendritic sites. The line represents a single exponential fit with length constant of 642 μm. **e,** Charging patterns of nucleated patches excised from human and rat L2/3 pyramidal neurons. Each symbol shows a sampling point, and the blue lines represent single exponential fits to the data. Photomicrographs of the recorded human and rat nucleated patches are inset. **f,** Specific membrane capacitance of human and rat L2/3 pyramidal neurons are not significantly different (n.s.d.), bars represent mean ± S.D. **g,** Simultaneous dendritic and somatic recordings of simulated EPSP trains, showing the pattern of D-S and S-D attenuation in a human and rat L2/3 pyramidal neuron. The driving currents injected at dendritic and somatic sites are shown below voltage traces. **h,** Somatic impact of the area of dendritic sEPSPs, calculated by the division of the somatic area of sEPSP trains generated at the soma and apical dendritic sites. The line represents a single exponential fit with a length constant of 679 μm. **i,** Somatic impact of dendritic sEPSP trains following normalization of the amplitude of the first sEPSP of the train in RL2/3 (open symbols) and HL2/3 (filled symbols) pyramidal neurons. Data have been fit by linear regression (rat, red line; human, blue line). The inset shows representative somatic recording of amplitude-normalized sEPSP trains, generated at somatic and apical dendritic sites. **j,** Somato-dendritic peak voltage attenuation of negative voltage steps in human and rat L2/3 pyramidal neurons. The line represents a single exponential fit with a length constant of 708 μm. The inset shows a representative recording of a HL2/3 pyramidal neuron.

As the attenuation and filtering of synaptic potentials in neurons is principally determined by the passive properties of the somatic and dendritic membrane ^38^, such distance-dependent relationships would be expected to diverge if biophysical properties such as specific membrane capacitance were distinct between species. Notably, recent work has shown that the specific membrane capacitance of HL2/3 pyramidal neurons is significantly lower than that of RL2/3 pyramidal neurons ^39^, and divergent from previous measurements ^40^, a biophysical adaptation predicted to enhance the somatic impact of dendritic EPSPs and accelerate their time-course ^39^. As our direct dendritic recordings do not support such a species-specific adaptation, we next determined the specific membrane capacitance of human and rat neurons by examining the charging responses of nucleated patches excised from identified L2/3 pyramidal neurons ^40^ (Fig. 2e). In each nucleated patch examined, charging responses evoked by brief, small amplitude voltage pulses (7 ms, −5 mV) were well described by a single exponential function, with time constants of 21.0 ± 1.8 μs in HL2/3, and 24.7 ± 1.6 μs in RL2/3 pyramidal neurons (Fig. 2e). Following measurement of the geometry of nucleated patches we calculated near identical values of specific membrane capacitance in human and rat L2/3 pyramidal neurons (human: 0.98 ± 0.03 μF/cm^2^; rat: 0.97 ± 0.04 μF/cm^2^; t-test, df*= 29, P=* 0.91; Fig. 2f). Together, therefore, these data reveal that the attenuation of single sEPSPs in the dendritic tree of human and rat L2/3 pyramidal neurons obeys a similar distance-dependent rule, governed by invariant biophysical constants. Given this, our findings indicate that the electrical architecture of the apical dendritic tree of L2/3 pyramidal neurons is not conserved between species, with the physically larger HL2/3 pyramidal neurons also exhibiting greater electrical distribution.

We next investigated if such relationships diverged when the complexity of simulated synaptic waveforms was increased. When trains of five closely spaced sEPSPs were generated at dendritic sites, we found that the distance-dependent decrement of the somatic impact of dendritic sEPSP trains fell on a similar curve in human and rat L2/3 pyramidal neurons (sEPSP trains generated at 50 Hz; driving current: amplitude= 200 pA; τ_rise_= 0.5 ms; τ_decay_= 5 ms; length constant= 679 μm; Fig. 2g,h). Notably, however, when the effects of dendritic cable filtering on the somatic amplitude of sEPSP trains was compensated for by scaling the amplitude of the first sEPSP of trains ^41^, a normalization of the somatic impact of sEPSP trains generated at dendritic and somatic sites was apparent in HL2/3 (slope= −0.14 per 1 mm), but not RL2/3 pyramidal neurons (slope= 0.6 per 1 mm; Fig. 2i), a phenomenon only previously observed in other neuronal classes in the rodent neocortex and hippocampus ^41,42^. Consistent with this, in HL2/3 pyramidal neurons the somatic half-width of single dendritic sEPSPs generated widely across the apical dendritic tree was not determined by the site of generation in the dendritic tree, but was similar to sEPSPs generated at the soma (slope: 0.15 per 1 mm; Supplementary Fig. 3). In other neuronal classes, such time-course normalization is mediated by the sculpting of EPSPs by the influence of hyperpolarization-activated cyclic nucleotide-modulated (HCN) channels ^41,42^. Consistent with the involvement of HCN channels in this behaviour we observed that HL2/3 pyramidal neurons exhibited a greater degree of time-dependent rectification during somatic hyperpolarizing voltage responses (Supplementary Fig. 4a), as has previously been described ^43^, and showed a pronounced crossover in the decay phase of sEPSPs simultaneously recorded at dendritic site of generation and the soma, a hallmark of non-uniform membrane conductance ^41,42^ (Fig. 2a, Supplementary Fig. 3a). Furthermore, the resting membrane potential of HL2/3 pyramidal neurons was found to be significantly depolarized with respect to RL2/3 pyramidal neurons (Supplementary Fig. 4b). The pharmacological blockade of HCN channels with ZD 7288 (10 μM) ^41,42^ powerfully hyperpolarized the resting membrane potential of HL2/3 pyramidal neurons, increased the apparent input resistance, and abolished the normalization of sEPSP time-course and summation (Supplementary Fig. 3). Taken together these findings indicate that a constraint of EPSP time-course and temporal summation, coupled with the larger physical length of the apical dendritic tree of HL2/3 pyramidal neurons, limits the direct impact that dendritic excitatory input has at the axonal site of AP generation ^44^, suggesting that apical dendritic excitatory synaptic input may be locally integrated, and signaled into neuronal output by the generation of dendritic electrogenesis.

### Phylogenetically conserved apical dendritic spike generation

As a first step to explore the electrical excitability of the dendritic tree of human and rat L2/3 pyramidal neurons we examined the properties of AP backpropagation (Fig. 3). Notably, the properties of somatically recorded APs were distinct between rat and human L2/3 pyramidal neurons, but were similar between HL2/3 pyramidal neurons recorded from tissues obtained from patients who had undergone neurosurgery for the alleviation of refractory epilepsy or the removal of sub-cortical brain tumours (Supplementary Fig. 5a-e). Simultaneous dendritic recordings revealed that somatic current step-evoked APs backpropagated into the apical dendritic tree (Fig. 3a,b). In RL2/3 pyramidal neurons, the amplitude of backpropagating APs (BPAPs) monotonically decreased, and the time-delay between somatic APs and BPAPs increased as dendritic recordings were made from increasingly distal sites (linear fit: conduction velocity= 0.21 m/s, r^2^= 0.90, n= 37; Fig. 3c,d, black open symbols, full line). In contrast, in HL2/3 pyramidal neurons the distance-dependent pattern of AP backpropagation was more heterogeneous (Fig. 3c). In distal apical dendritic recordings, a fraction of HL2/3 pyramidal neurons exhibited BPAPs that appeared to follow the distance-dependent pattern of BPAP attenuation recorded from RL2/3 pyramidal neurons, whereas others exhibited larger amplitude BPAPs (Fig. 3c, black symbols, extrapolated fit to rat data shown as a dashed line). This heterogenous pattern of AP backpropagation was unrelated to the properties of somatically recorded APs, or etiology, as both weak and strong BPAPs were observed in neurons obtained from epileptic or tumour resection surgeries (Supplementary Fig. 5f,g). Despite this heterogeneity, APs backpropagated into the apical dendritic tree of HL2/3 pyramidal neurons with a conduction velocity of 0.23 m/s, similar to that of RL2/3 pyramidal neurons (linear fit, r^2^= 0.87, n= 65; Figure 3a,d, filled black symbols).

**Figure 3.**
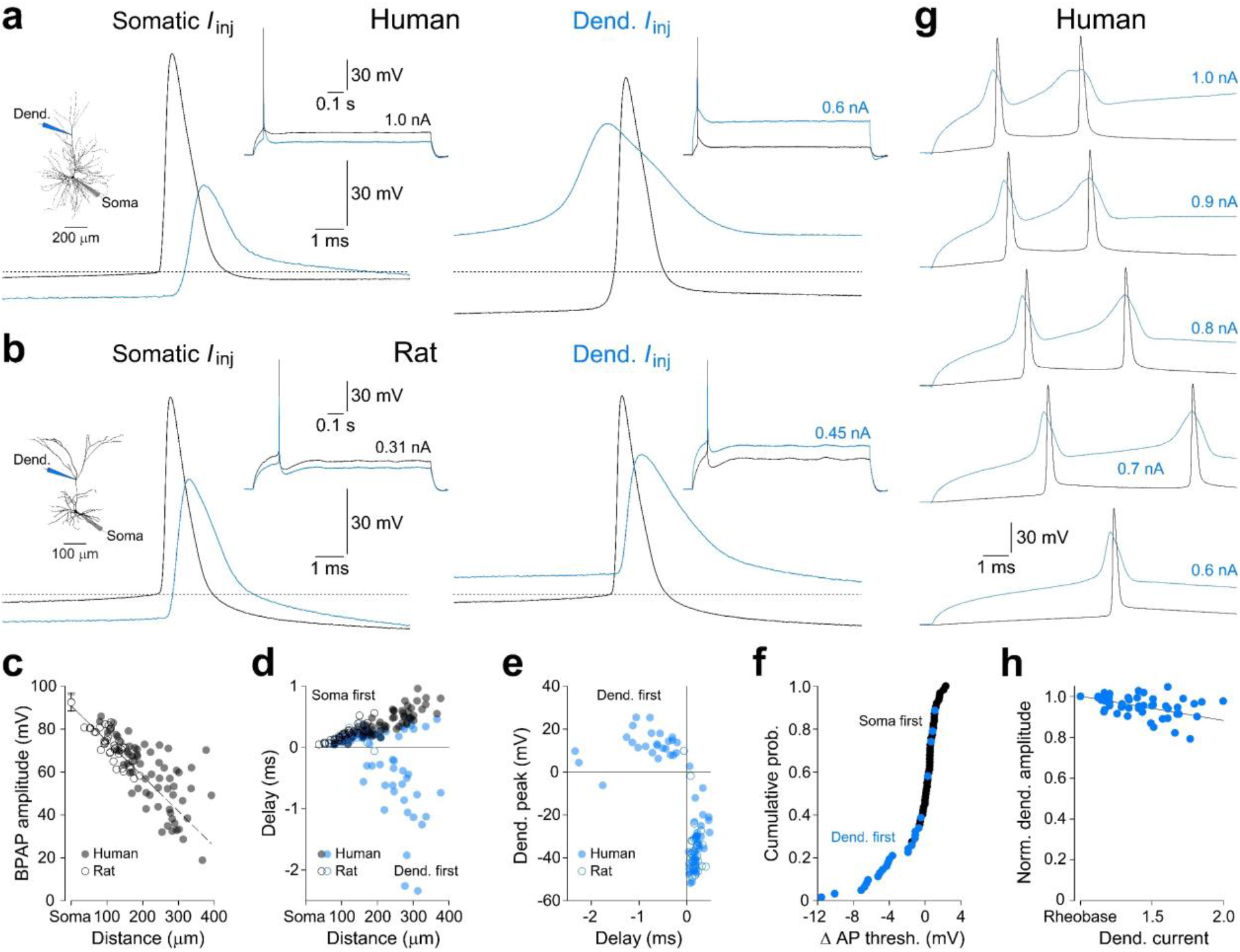
Dendritic excitability of human and rat L2/3 pyramidal neurons. **a,** Dendritic spike generation in response to the presentation of threshold dendritic positive current step (blue trace, Dend I_inj_) in a HL2/3 pyramidal neuron. Note that the dendritic spike forward propagates to drive AP firing which has a hyperpolarized threshold (dashed line). The morphology of the recorded neuron and full evolution of voltage responses are shown inset, together with the amplitude of current steps. **b,** Threshold positive current steps delivered at somatic and dendritic sites in a RL2/3 pyramidal neuron evoke similar patterns of regenerative activity. **c,** Quantification of the distance-dependency of BPAP amplitude in rat (open symbols) and human (filled symbols) L2/3 pyramidal neurons. The average amplitude (± S.D.) of APs at the soma are shown. The continuous line represents a fit to the distance-dependent decrement of BPAPs in RL2/3 pyramidal neurons and has been extrapolated (dashed line). **d,** The delay between somatic APs and BPAPs (black symbols) generated by somatic I_inj_ and regenerative spikes evoked by dendritic I_inj_ (blue symbols). Note the distance-dependent generation of dendritic spikes which precede APs (negative times) in HL2/3 pyramidal neurons. **e,** At the time of AP initiation the voltage of dendritic spikes which preceded AP firing (negative times) typically overshot 0 mV in HL2/3 pyramidal neurons (filled symbols). **f,** Cumulative probability distribution of the difference in the voltage threshold for AP initiation when APs were driven by somatic and dendritic I_inj_ in HL2/3 pyramidal neurons. Blue symbols represent APs driven by dendritic spikes (Dend. first). **g,** Dendritic spikes are generated across a wide suprathreshold range in HL2/3 pyramidal neurons. **h,** The peak voltage of dendritic spikes does not markedly attenuate with increasing excitatory drive. The peak voltages of dendritic spikes are shown relative to threshold values (rheobase), to allow comparison between neurons; the line represents a linear regression.

Next, we directly examined apical dendritic electrical excitability by injecting incremental series of positive current steps through dendritic recording electrodes until a threshold (rheobase) regenerative response was generated. In recordings made from proximal apical sites, the injection of rheobase dendritic positive current steps directly drove AP firing in human and rat L2/3 pyramidal neurons (<150 μm from the soma; current step duration= 1 s; Fig. 3, positive times indicate that APs preceded BPAPs). In contrast, when recordings were made from more distal sites of HL2/3 pyramidal neurons, dendritic positive current steps typically evoked, as a threshold response, all-or-none dendritic spikes, which overshot 0 mV and preceded and drove AP firing following their forward propagation to the soma and axon (Fig. 3, negative times indicate dendritic spikes preceding APs, the amplitude of dendritic events was measured at the time of AP initiation). In dendritic recordings made at sites > 200 μm from the soma, dendritic spikes were generated as a threshold response in 22 of 35 HL2/3 pyramidal neurons examined (Fig. 3d). In these neurons the dendritic current required to initiate dendritic spikes, and consequentially AP output, was only marginally greater than the rheobase current required to directly generate AP firing when injected at the soma (dendrite rheobase: 0.56 ± 0.03 nA; soma rheobase= 0.50 ± 0.03 nA; distance from soma: 276 ± 9 μm). Notably, when AP firing was driven by dendritic spikes, somatically recorded APs were initiated from relatively negative membrane potentials, with a threshold that was significantly hyperpolarized in comparison to those evoked by somatic excitatory input (t-test, df= 24*, P*< 0.0001; Fig. 3a,f). Furthermore, dendritic spike generation in HL2/3 pyramidal neurons was sustainable across a wide range of dendritic current steps, and reliably drove AP output when the magnitude of dendritic current steps was increased to values up to 2 times the threshold current intensity (Fig. 3g,h). In contrast, in RL2/3 pyramidal neurons, the presentation of long threshold steps of dendritic current evoked dendritic spike initiation in only one recording (Fig. 3d). Taken together these data reveal that the apical dendrites of HL2/3 pyramidal neurons are highly electrically excitable, and capable of generating powerful dendritic spikes. Our findings do not, however, directly show that apical dendritic excitability is enhanced in HL2/3 pyramidal neurons, as central neurons are not physiologically driven by long steps of current. We therefore explored if more physiological EPSP-shaped patterns of dendritic excitatory input ^10^ led to the initiation of dendritic spikes in human and rat L2/3 pyramidal neurons.

To our surprise we found that dendritic spikes were generated in response to threshold EPSP-shaped current waveforms in both HL2/3 and RL2/3 pyramidal neurons (τ_rise_= 0.5 ms; τ_decay_= 5 ms; Figs. 4 and 5). The presentation of single threshold sEPSPs, even at relatively proximal dendritic sites, typically led to the initiation of stereotyped dendritic spikes in human and rat L2/3 pyramidal neurons, which forward propagated to the soma and axon to drive AP firing (Fig. 4a-d). Analysis demonstrated that at apical dendritic sites >100 μm from the soma, dendritic spikes typically led and drove AP firing in both species (Fig. 4e), to produce a timing relationship of regenerative activity that was distinct from the relationship evoked by threshold somatic sEPSPs, which simply reflected the progressive time delay between APs and dendritic BPAPs (Fig. 4e). In common with dendritic spikes generated in response to long current steps, the voltage threshold for AP initiation, recorded at the soma, was significantly hyperpolarized when APs were driven by dendritic spikes in both human and rat L2/3 pyramidal neurons (human: dendritic EPSP driven= −62.8 ± 1.1 mV, soma EPSP driven= −50.6 ± 0.6 mV, t-test, df= 22, *P*< 0.0001; rat: dendritic EPSP driven= −60.1 ± 1.0 mV, soma EPSP driven= −52.3 ± 0.8 mV, t-test, df= 15, *P*< 0.0001; Fig. 4f). To examine the factors underlying this, we calculated the first derivative of somatically recorded APs, finding that dendritic spike initiation led to the generation of a ramp-like component which preceded the rapid upstroke of the first derivative of APs, a component that was absent when APs were driven by somatic sEPSPs in human and rat L2/3 pyramidal neurons, and so reflects the somatic impact of forward propagating dendritic spikes (Fig. 4g). Notably, the generation of dendritic spikes by sEPSPs was not simply a threshold mechanism, but was sustainable across a wide range of suprathreshold excitatory input, with grouped data showing a flat relationship between the peak voltage of dendritic spikes and the amplitude of the underlying simulated synaptic current in human and rat L2/3 pyramidal neurons (Fig. 4h), an input range over which dendritic spikes reliably preceded and drove AP output (Fig. 4i). These data therefore demonstrate that apical dendritic spike generation in human and rat L2/3 pyramidal neurons does not exhibit anti-coincidence properties.

**Figure 4.**
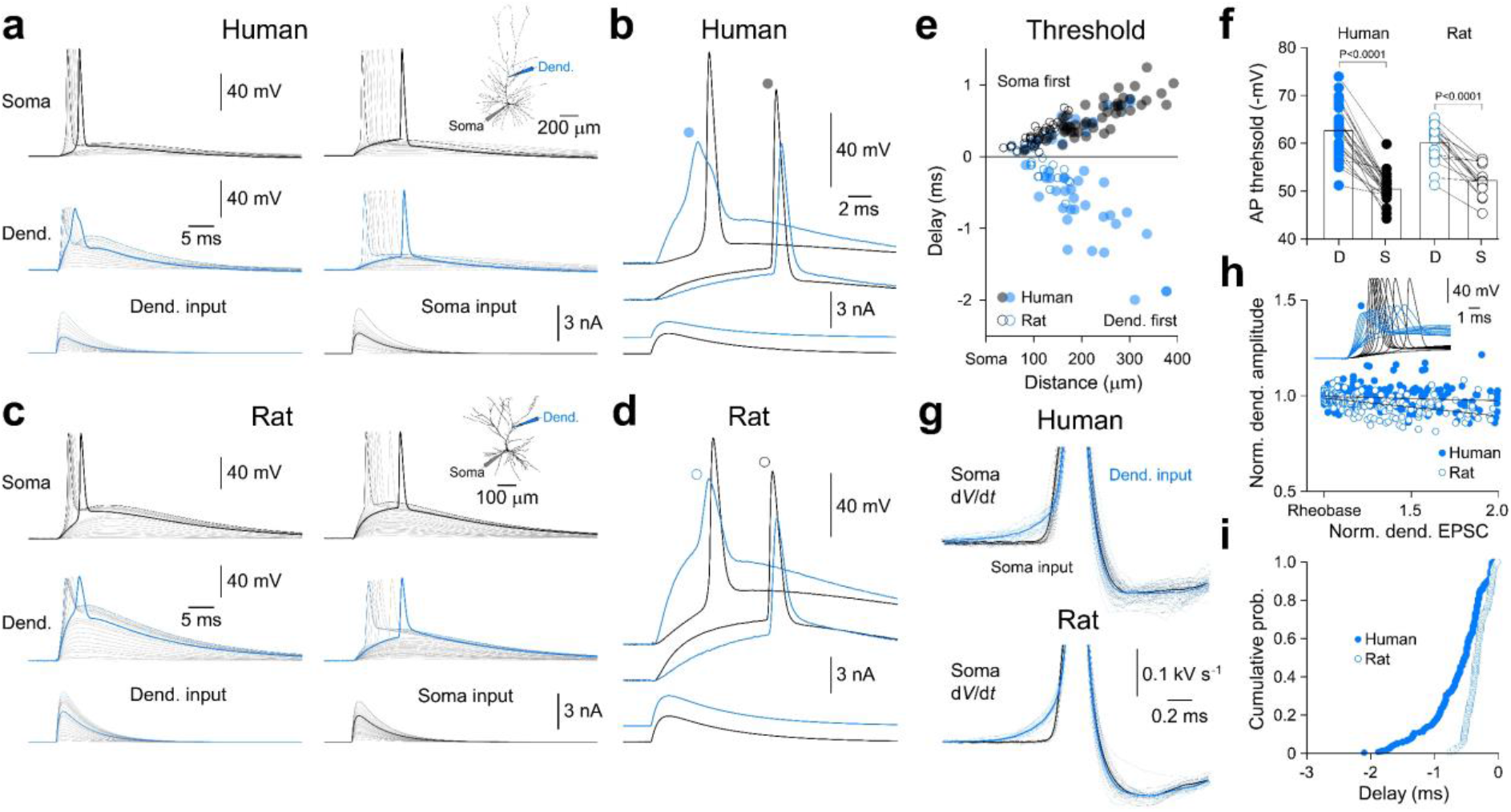
Stereotyped dendritic spikes are driven by physiologically relevant stimuli in human and rat layer 2/3 pyramidal neurons. **a,** Dendritic spikes (Dend, blue) drive AP firing in HL2/3 pyramidal neurons when generated by sEPSPs. Note that families of dendritic sEPSPs evoke dendritic spikes at threshold (demarked by bold traces), a pattern of activity preserved as the amplitude of the sEPSPs is further increased. In contrast, families of somatic sEPSPs evoke APs, followed by dendritic BPAPs. The morphology of the neuron and placement of recording electrodes are shown inset. **b,** Threshold firing patterns evoked by dendritic and somatic sEPSCs at a higher time-base. **c,d,** Near identical pattern of active dendritic integration of simulated synaptic input in a RL2/3 pyramidal neuron. **e,** Pooled data show that threshold somatic sEPSPs generated APs which preceded dendritic BPAPs (black symbols). In contrast, threshold sEPSPs generated at dendritic sites >100 μm from the soma typically evoked dendritic spikes which preceded APs (negative times) in HL2/3 (filled blue symbols) and RL2/3 (open blue symbols) pyramidal neurons. **f,** Significant difference in AP initiation threshold when AP firing was evoked by dendritic (D) sEPSPs which drove dendritic spikes, and somatic (S) sEPSCs (t-test, *P* values shown). **g,** First derivative (d*V*/d*t*) of somatically recorded APs evoked by dendritic (blue) and somatic input (black) in HL2/3 (27 overlain recordings) and RL2/3 (17 overlain recordings) pyramidal neurons. The bold traces represent digital averages; note the characteristic ramp preceding the AP upstroke when APs were driven by dendritic spikes (blue traces, Dend. input). **h,** Dendritic spike amplitude does not decrement when generated across a wide suprathreshold sEPSC range in HL2/3 (filled symbols) and RL2/3 (open symbols) pyramidal neurons. Values are relative to threshold (rheobase) and have been fit by linear regression. The inset shows overlain dendritic (blue, 260 μm from soma) and somatic voltage responses evoked by sEPSP families in a human L2/3 pyramidal neuron. **i,** Cumulative probability distribution of time delay between dendritic spikes and APs for suprathreshold events generated by families of dendritic sEPSPs shown in panel h.

**Figure 5.**
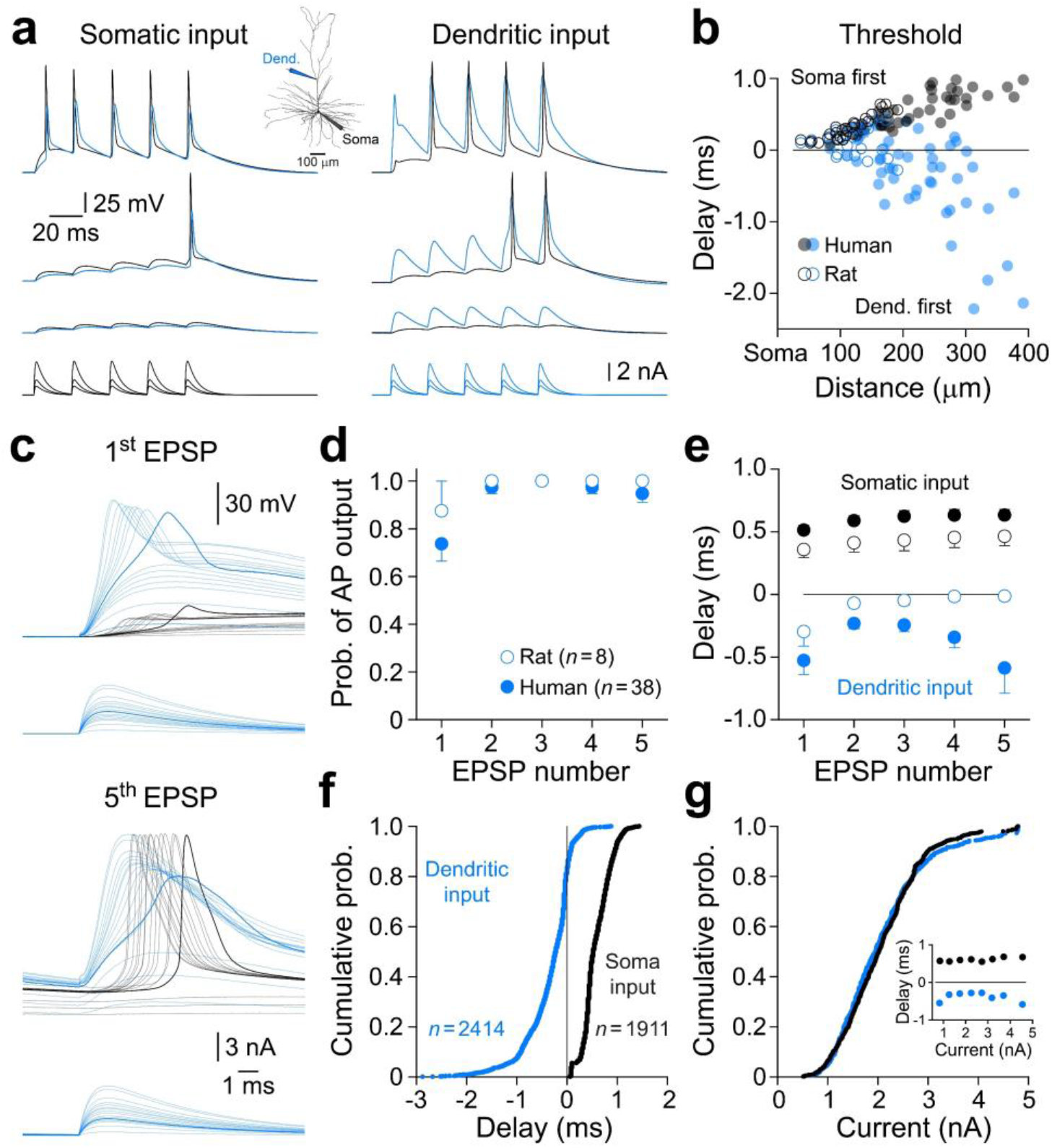
Dendritic spikes can be generated at high frequency in human and rat L2/3 pyramidal neurons. **a,** Repetitive dendritic spike generation driven by simulated EPSP trains (lower overlain traces) in HL2/3 pyramidal neurons. The morphology of the neuron and placement of recording electrodes is shown inset. **b,** Timing relationship between regenerative events evoked by threshold sEPSP trains generated at the soma (black symbols) or dendrite (blue symbols). **c,** Baselined first and fifth sEPSPs of the train at a faster time base (same neuron as A). Note that the first sEPSP is crowned by the generation of dendritic spikes which drive somatic spikelets across a wide suprathreshold range, while the last sEPSP of the train is crowned by dendritic spikes which drive AP output. Threshold events are shown in bold. **d,** Pooled data describing the probability of AP firing generated in response to suprathreshold sEPSP trains in human and rat L2/3 pyramidal neurons. **e,** Pooled data showing the timing of regenerative events for each sEPSP of the train. **f,** Cumulative probability distribution of the timing of all regenerative events evoked by dendritic (blue) and somatic (black) sEPSP trains in HL2/3 pyramidal neurons. Note the clear disparity between the timing relationship and the preponderance of APs driven by dendritic spikes (negative times) when sEPSP trains were presented at dendritic sites. **g,** Cumulative probability distributions of amplitude of the current underlying sEPSPs (dendritic, blue), which drove the regenerative events shown in (f). The inset shows the average time delay between the regenerative events evoked by somatic and dendritic EPSPs at the indicated binned amplitudes of the driving current.

We next explored if dendritic spikes could be generated repetitively. To do so we presented trains of sEPSPs at dendritic and somatic recording sites (inter-event interval 20 ms; Fig. 5). In both human and rat L2/3 pyramidal neurons the injection of threshold sEPSP trains at dendritic sites >100 μm from the soma led to the initiation of dendritic spikes, which were characteristically generated late in the sEPSP train (Fig. 5a, Supplementary Fig. 5). At threshold, dendritic spikes forward propagated to drive AP output, yielding a distance-dependent timing relationship distinct from that generated by the generation of threshold sEPSP trains at the soma (Fig. 5b, Supplementary Fig. 6). Remarkably, when the amplitude of excitatory input was increased, dendritic spikes could be generated at high frequencies (Fig. 5a). Analysis revealed that, in the majority of human and rat L2/3 pyramidal neurons, dendritic spikes crowned the peak of each sEPSP of the train, and subsequently drove AP firing (Fig. 5a,d,e, Supplementary Fig. 6). In a fraction (<25 %) of HL2/3 pyramidal neurons, however, dendritic spike generation in response to the first sEPSP of a train failed to drive AP output, but was detectable at the soma as a subthreshold spikelet (Fig. 5a,c). This suppression of neuronal output was, however, transient, as dendritic spikes generated later in sEPSP trains reliably forward propagated to initiate AP output (Figure 5a,b,d). Indeed, pooled data revealed that active dendritic synaptic integration of sEPSP trains was operational across a wide input range, with dendritic spikes evoked at current intensities up to 4-fold greater than threshold (Fig. 5f,g). Over this input range, 2021 of 2414 APs were driven by dendritic spikes, with grouped data showing a clear difference from the timing of regenerative activity evoked by the presentation of somatic sEPSP trains over a similar range (Fig. 5f,g). Taken together therefore, these findings demonstrate that apical dendritic spikes are reliably and repetitively generated across a wide range of excitatory input, suggesting that the dendritic integrative properties of both human and rat L2/3 pyramidal neurons efficiently implement a correlative computation.

To further examine the frequency-following capacity of dendritic spikes in L2/3 pyramidal neurons we injected series of short (3 ms) threshold positive current steps at distal dendritic recording sites (Fig. 6). In both human and rat L2/3 pyramidal neurons dendritic spikes could be initiated and drive AP firing in response to dendritic positive current steps delivered at intervals as short as 10 ms (Fig. 6a,b). This frequency-following capacity mirrored the generation of AP firing evoked in response to short somatic positive current steps (Fig. 6a,b), suggesting that dendritic spikes and APs are mediated by similar classes of voltage-gated ion channels. To test this idea, we locally and transiently applied the sodium channel blocker tetrodotoxin (TTX) to a site close to the dendritic recording electrode (2 μM; pipette separation <20 μm; Fig. 6e,f). Under control conditions dendritic spikes robustly forward propagated to drive AP firing in HL2/3 pyramidal neurons; however, following the local dendritic application of TTX, dendritic spike generation was eliminated, and could not be restored by increasing the amplitude of positive current steps (Fig. 6e). This effect of dendritic sodium channel blockade was fully reversible and confined to dendritic sites, as the somatic amplitude of APs generated in response to short somatic current steps was unperturbed (Fig. 6e,f). Furthermore, the bath application of TTX (0.5-1 μM) abolished dendritic spikes evoked by short (3-5 ms) and long (1 s) positive dendritic current steps, as well as single and trains of sEPSPs (Supplementary Fig. 7a-d), revealing, by subtraction, that dendritic spikes in HL2/3 pyramidal neurons had an amplitude of 40.7 ± 2.4 mV and exhibited fast rise and decay kinetics (10-90 % rise time= 0.70 ± 0.07 ms; half-width= 1.51 ± 0.16 ms; n= 14; Supplementary Fig. 7d). Similarly, in RL2/3 pyramidal neurons, dendritic spikes were found to be mediated by dendritic TTX-sensitive sodium channels and possessed comparable properties (amplitude= 52.6 ± 2.2 mV; 10-90 % rise-time= 0.53 ± 0.04 ms; half-width= 1.46 ± 0.10 ms; n= 7; Supplementary Fig. 7d-f). Taken together, these data reveal that dendritic spikes in human and rat L2/3 pyramidal neurons are mediated by the regenerative activation of dendritic sodium channels.

**Fig. 6.**
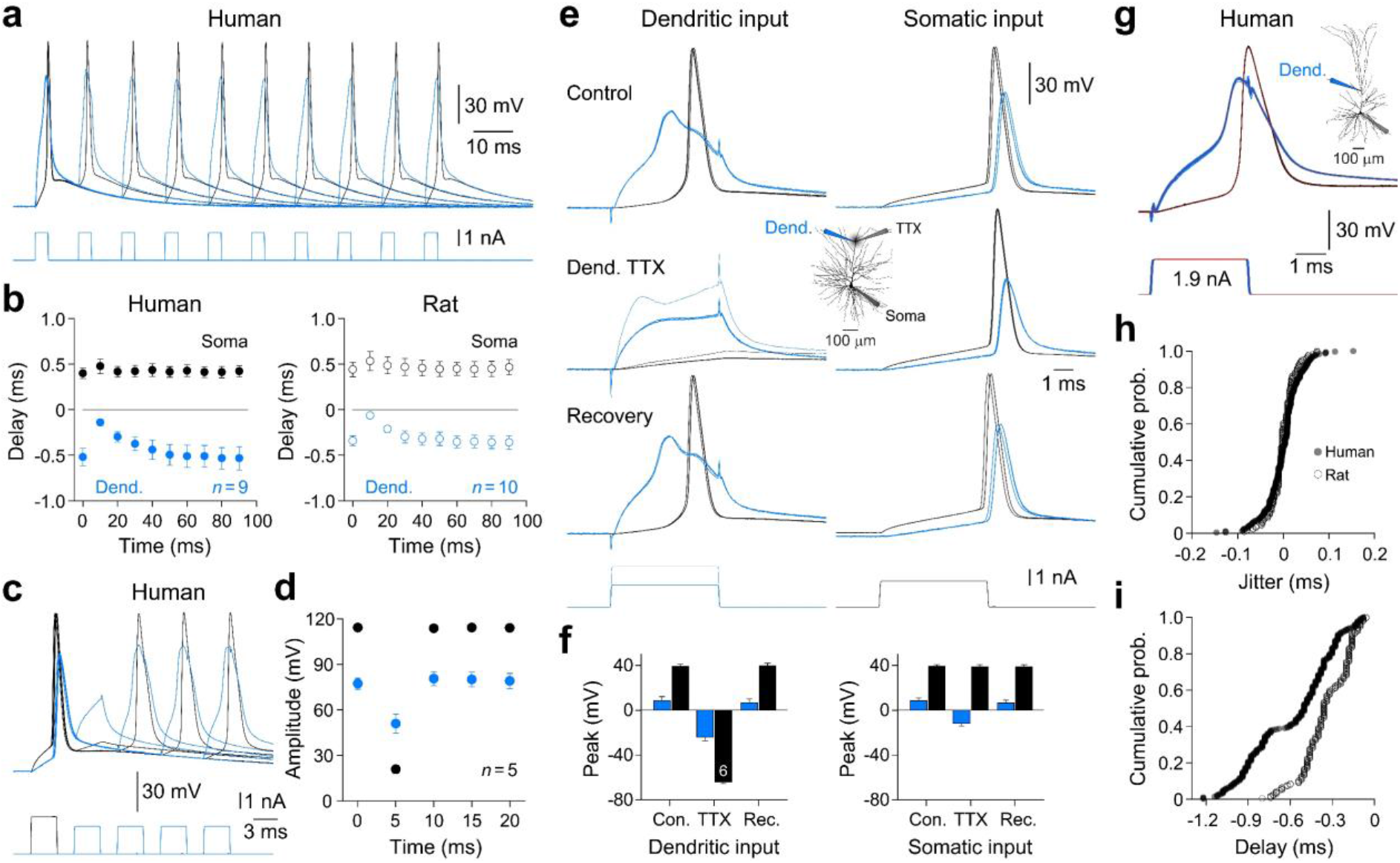
High-fidelity dendritic spikes are mediated by dendritic sodium channels in human L2/3 pyramidal neurons. **a,** Initiation of dendritic spikes at high repetition frequencies in HL2/3 pyramidal neurons. Each dendritic spike was generated in response to a short positive current step (steps separated by Δ10 ms). **b,** Quantification showing that dendritic spike generation leads AP firing (negative times) in human and RL2/3 pyramidal neurons when generated by dendritic current steps (Dend.) at the indicated intervals. **c,** AP initiation transiently precludes dendritic spike generation. In each overlain trial, AP firing was driven by the presentation of a threshold somatic current step (black, lower trace), which was followed in time by the presentation of threshold dendritic current steps at Δ5 ms intervals; note that at the first interval dendritic spike initiation did not occur. **d,** Pooled data of the peak amplitude of somatic (black) and dendritic (blue) voltage responses showing the transient suppression of dendritic spike generation by AP firing. **e,** Dendritic spikes are mediated by sodium channels in HL2/3 pyramidal neurons. Overlain voltage traces show dendritic (blue) and somatic (black) voltage responses evoked by brief dendritic or somatic positive current steps (lower traces). The transient local dendritic application of TTX (2 μM) blocked the initiation of dendritic spikes but did not alter the characteristics of APs generated by somatic positive current pulses. A partial reconstruction of the morphology of the HL2/3 pyramidal neuron is shown inset, illustrating the recording and TTX application sites. **f,** Pooled data describing the blockade of dendritic spikes and the reduction of the amplitude of BPAPs by the local dendritic application of TTX. Bars resent mean ± S.E.M.. **g,** Overlain voltage traces (30 trials) aligned at the peak of somatic APs show the minimal trial-to-trial variation in dendritic spike to AP coupling around digital averages (red traces). The morphology of the HL2/3 pyramidal neuron is shown inset. **h,** Trial-to-trial temporal jitter around the mean dendritic spike to AP time-delay for HL2/3 (filled symbols, n= 12) and RL2/3 (open symbols, n= 11) pyramidal neurons. **i,** Cumulative probability distribution of the time-delay between dendritic spikes and APs for the data shown in panel h.

To explore if dendritic excitatory input to human and rat L2/3 pyramidal neurons could evoke regenerative dendritic activity following the pharmacological blockade of sodium channels, we generated families of large amplitude sEPSPs, and positive voltage steps, in the presence of TTX (Supplementary Figs. 7c and 8). Under these conditions dendritic excitatory input did not evoke regenerative activity, despite the generation of intense dendritic depolarization (Supplementary Figs. 7c and 8). We noted, however, that such input-output relationships were highly rectified in both species, suggesting that voltage-gated potassium channels regulate dendritic electrical excitability (Supplementary Fig. 8 a-c, f,g). Consistent with this idea, bath application of the potassium channel blocker 4-amino pyridine (5 mM), in TTX, was found to unmask apical dendritic electrogenesis generated in response to dendritic sEPSPs or positive current steps, which was largest in amplitude and peaked first at dendritic recording sites, and decrementally spread to the soma (Supplementary Fig. 8a-g). This dendritic regenerative activity could be pharmacologically blocked by the application of the non-specific calcium channel antagonist cadmium (100 μM, n= 5 HL2/3 pyramidal neurons; Supplementary Fig. 8d,e). Notably, dendritic calcium spikes in HL2/3 pyramidal neurons recorded under these conditions discharged repetitively throughout the course of long dendritic positive current steps, and exhibited amplitudes similar to the dendritic calcium APs previously reported for HL2/3 pyramidal neurons under control conditions ^22^ (cadmium subtracted dendritic calcium spike: dendrite= 30.9 ± 6.6 mV (range= 24.8 - 40.4 mV), soma= 23.4 ± 4.2 mV n= 5 neurons). Therefore, consistent with observations made from other neuronal classes ^25,45^, our findings demonstrate that the expression of dendritic calcium electrogenesis is powerfully controlled by the activation of voltage-gated potassium channels in human and rat L2/3 pyramidal neurons.

In other central neurons a commonality between the generation mechanisms of APs and dendritic spikes has been shown to limit the temporal dynamics of active dendritic integration, as reliance on the same classes of ion channel can lead to use-dependent inactivation ^46^. To explore the time-window over which AP output influences dendritic sodium spike generation in HL2/3 pyramidal neurons we generated AP firing by the presentation of short somatic current pulses, and at subsequent times probed for the generation of dendritic spikes by the delivery of threshold dendritic positive current steps (Fig. 6c,d). To our surprise AP firing and subsequent AP backpropagation only precluded dendritic integration over a very narrow time-window (<10 ms), after which dendritic spikes were reliably generated and forward propagated to drive AP output (Fig. 6c,d). Thus, in human and rat L2/3 pyramidal neurons dendritic spikes signal, at high repetition frequencies, the result of dendritic integration, but how precisely do dendritic spikes drive neuronal output? To address this, we generated dendritic spikes in response to short threshold positive current steps and measured the time-delay between the peak of dendritic spikes and somatically recorded APs across multiple trials (Fig. 6g,h). Pooled data revealed that the forward propagation of dendritic spikes was reliable and drove time-locked AP firing, exhibiting little trial-to-trial temporal jitter (half-width of the temporal jitter around mean: HL2/3= 61 μs; RL2/3= 41 μs; Fig. 6g,i). This temporal precision emerged in HL2/3 pyramidal neurons despite the relatively long time-delay incurred by the forward propagation of dendritic spikes from the site of generation to the axon (Fig. 6i). Taken together, therefore, our findings reveal that a phylogenetically conserved mechanism signals the result of dendritic computations into AP output in human and rat L2/3 pyramidal neurons.

## Discussion

Here we demonstrate the mechanistic and functional conservation of the properties of active dendritic integration in human and rodent L2/3 neocortical pyramidal neurons. Our findings do not, however, support conservative scaling principles, which posit that the electrical architecture of neuronal classes is conserved between species ^47^. This arises because voltage attenuation in the apical dendritic tree of L2/3 pyramidal neurons was found to exhibit similar distance-dependent properties between species, and so the electrical architecture of HL2/3 pyramidal neurons is more distributed, reflecting the greater physical length of apical dendritic tree. Such a finding is noteworthy, as previous computational modeling has suggested that the dendritic cable properties of human L2/3 pyramidal neurons are distinct from those of the rodent, due to unique capacitive properties of the plasma membrane ^39^. However, our simultaneous recordings, and direct measurement of membrane capacitance do not support such a species-specific adaptation, a finding underscored by observations made from HL5 pyramidal neurons ^24^. In contrast, we find that the electrical and morphological distribution of the dendritic tree provides a foundation for the enhanced emergence of independent integrative compartments in HL2/3 pyramidal neurons. At these sites, dendritic excitatory input is integrated to form dendritic sodium spikes, which forward propagate to drive neuronal output across a wide range of correlated excitatory input, consistent with computational models which describe the dendritic integrative operations of rodent pyramidal neurons as two-layered neural networks 9,32,33. As neuronal modeling has suggested that dendritic sodium spikes of rodent L2/3 pyramidal neurons are driven by the strongest 1% of excitatory synaptic inputs, and by the activation of spatially clustered excitatory input ^28^, it is tempting to speculate that the increased number of dendritic spines in HL2/3 pyramidal neurons ^3,5,6^ provides an increased substrate for such dendritic computations.

In both human and rat L2/3 pyramidal neurons, dendritic sodium spikes could be generated at high frequencies, exhibiting characteristics similar to those recorded in isolation from the thin apical dendrites of rodent neocortical pyramidal neurons *in vivo* using invasive and non-invasive recording techniques ^14,19^, and other neuronal classes ^30,35,36^. However, these properties are mechanistically and functionally distinct from the pattern of apical dendritic excitability recently reported for HL2/3 pyramidal neurons, characterized by the generation of calcium channel-mediated dendritic APs ^22^, which inactivate upon increased excitatory drive to implement a novel anti-coincidence computation. Such a profound difference in the active dendritic integrative operations of HL2/3 pyramidal neurons cannot be ascribed to methodological differences between studies, as recordings were made under similar ionic conditions from the same cortical area. It is therefore possible that through the evolutionary expansion of the human neocortex, neuronal classes with distinct dendritic integrative properties have emerged, as highlighted by the existence of a sparse population of molecularly unique layer 3 pyramidal neurons in the human ^4^, which execute complimentary computations. On the other hand, our observation that the pharmacological blockade of voltage-gated potassium channels unmasks dendritic calcium electrogenesis in human and rat L2/3 pyramidal neurons, consistent with previous findings made from other neuronal classes in the rodent neocortex ^25,45^, suggests that differences between our and previous work ^22^ may, in part, be explained by the differential regulation of potassium channel function. Consistent with this idea, the membrane expression and voltage-dependent availability of voltage-gated potassium channels are known to be controlled by a variety of factors such as neuronal activity ^48–50^, osmolarity ^51^, intracellular calcium ^52^, and neuromodulators ^53^, and so may have been down-regulated in previous investigation of HL2/3 pyramidal neurons ^22^. This interpretation may help to explain the narrow range over which excitatory input could generate dendritic calcium spikes in previous investigation of HL2/3 pyramidal neurons ^22^. Notwithstanding, our observation of the conservation of suprathreshold dendritic integrative operations, biophysical constants, and dendritic cable properties between L2/3 pyramidal neurons of the human and rodent neocortex is consistent with the predominant molecular conservation of supragranular pyramidal neurons in these species ^4^. Thus, we suggest that the enlargement and complexification of the dendritic tree of human L2/3 pyramidal neurons provides an enhanced framework for the execution of phylogenetically conserved dendritic computational operations, which are translated into neuronal output by the high-fidelity generation and forward propagation of dendritic sodium spikes.

## Methods

### Human brain slice preparation

Males or females aged 17-70 years (median 42 years) who provided informed written consent for the provision of normative tissue surplus to diagnostic requirements (normative tissue) were admitted to the study. All clinical, experimental, and administrative procedures were approved by the Mater Misericordiae Ltd. Human Research Ethics Committee (HREC/MML/63124) and ratified by the The University of Queensland. Patients undergoing neurosurgery were maintained under general anesthesia following neuro-muscular blockade (Supplementary Tables 1 and 2). Brain tissue was resected typically less than 1 h after the start of the procedure and was considered surplus normative tissue. Normative tissue did not demonstrate abnormalities at the level of preoperative magnetic resonance imaging (MRI) or during neurosurgical resection and was not subjected to clinical pathological analysis. In refractory epilepsy cases, the epileptogenic zone (EZ) was identified by anatomoelectroclinical correlations. If there was discordance in these data, the EZ was localized by stereo-electroencephalography prior to resection (see Supplementary Table 1). Normative tissue adjacent to the EZ was resected, while tissue obtained from the EZ was subsequently resected and subjected to clinical pathological analysis (Supplementary Table 1). In tumour cases, the location of sub-cortical tumours was visualized by preoperative MRI and in some cases dynamically during neurosurgery with the use of optical imaging agents (Supplementary Table 2). In all cases the resected tissue provided was considered normative tissue during neurosurgery, whereas abnormal tissue was subjected to clinical pathological analysis (Supplementary Table 2). In all reported cases brain tissue was resected from the anterior temporal lobe (see Supplementary Tables 1 and 2). The resected brain tissue was immediately (<15 s) placed into a container and immersed in a saturated (95% O_2_, 5% CO_2_) ice-cold solution of the following composition: (in mM) sucrose 75, NaCl 87, KCl 2.5, NaH_2_PO_4_ 1.25, NaHCO_3_ 25, glucose 10, Na pyruvic acid 3, MgCl_2_ 3, CaCl_2_ 0.5. The container was sealed, packed into a second vessel and placed on ice for transportation to the laboratory, where the tissue was removed and placed in fresh ice-cold saturated (95% O_2_, 5% CO_2_) solution of the same composition within 14-30 min of resection. The tissue block was then inspected, divided if necessary, and re-blocked perpendicular to the cortical surface. We made no attempt to remove the remaining meninges from the surface of tissue blocks as preliminary evidence suggested that such mechanical disturbance impacted the morphology of the fine-caliber apical tuft dendrites of pyramidal neurons. Tissue blocks were adhered to the stage of a vibrating blade microtome and rapidly immersed in the same ice-cold solution after which 300-350 μm brain slices were prepared. Each brain slice was then immediately transferred to a storage chamber filled with a solution of composition (in mM): NaCl 92, KCl 2.5, NaH_2_PO_4_ 1.2, NaHCO_3_ 30, HEPES 20, glucose 20, Na-L-ascorbate 5, Na pyruvic acid 3, thiourea 2, MgCl_2_ 10, CaCl_2_ 0.5, saturated with 95% O_2_, 5% CO_2_ at 34-36°C for 30 min. Brain slices were then individually transferred to storage chambers filled with a saturated (95% O_2_, 5% CO_2_) solution of composition (in mM): NaCl 125, KCl 2.5, NaH_2_PO_4_ 1.25, NaHCO_3_ 25, glucose 25, Na pyruvic acid 3, MgCl_2_ 6, CaCl_2_ 1 and maintained at room temperature (20-24 °C).

### Rat brain slice preparation

All experiments were approved by the Animal Ethics Committee of The University of Queensland in accordance with the Australian Code of Practice for the Care and Use of Animals for Scientific Purposes. Wistar rats (male, P30 - P45) were deeply anesthetized by inhalation of isoflurane and decapitated. Brain slices (300 μm) including the temporal associative area, with reference to ^54^ were cut in the coronal plane in an ice-cold solution of composition (in mM): sucrose 75, NaCl 87, KCl 2.5, NaH_2_PO_4_ 1.25, NaHCO_3_ 25, glucose 10, Na pyruvic acid 3, MgCl_2_ 3, CaCl_2_ 0.5. Brain slices were then immediately transferred to a storage chamber filled with a saturated (95% O_2_, 5% CO_2_) solution of composition (in mM): NaCl 125, KCl 2.5, NaH_2_PO_4_ 1.25, NaHCO_3_ 25, glucose 25, Na pyruvic acid 3, MgCl_2_ 6, CaCl_2_ 1 at 34-34°C for 30 min, and maintained at room temperature.

### Electrophysiological recordings

Individual brain slices were placed in a chamber perfused with a constant flow of artificial cerebral spinal fluid (ACSF) of composition (in mM): NaCl 125, NaHCO_3_ 25, KCl 2.5, NaH_2_PO_4_ 1.25, 1 CaCl_2_ 1, MgCl_2_ 1, Na pyruvic acid 3 and glucose 25 saturated with 95 % O_2_ and 5 % CO_2_ at 35-36.5 °C. In the majority of experiments, spontaneous synaptic activity was blocked by the addition of CNQX (10 μM, Tocris), DL-AP5 (50 μM, Tocris), SR 95531 (5 μM, Tocris), and CGP52432 (10 μM, Tocris). The ion channel blockers ZD7288 (10 μM, Tocris), cadmium chloride (100 μM, Merck), 4-amino pyridine (5 mM, Merck) and TTX (0.5-1 μM, Alomone) were added to the extracellular solution when indicated. In some experiments TTX (2 μM, dissolved in saturated ACSF) was applied close to the site (<20 μm) of dendritic recording by pressure application (20 to 30 s) from a pipette with characteristics similar to those used for dendritic whole-cell recordings. Recordings were made from pyramidal neurons identified by their soma shape and ascending apical dendrite projecting to layer 1, when visualized with video-enhanced infra-red differential interference contrast microscopy. Dual whole-cell, current-clamp recordings were made with identical dedicated current-clamp amplifiers (BVC 700A; Dagan). The electrode capacitance and series resistance were compensated using the active circuitry of the amplifier. Pipettes were filled with a solution containing (in mM): K-gluconate 135, NaCl 7, HEPES 10, Na2ATP 2, NaGTP 0.3, MgCl 2, Na2Phosphocreatine 10, and Alexa Fluor 598 0.01, pH 7.2 - 7.3, for somatic (open tip resistance 3-6 MΩ) and dendritic (open tip resistance 10-15 MΩ) recordings. Somatic or dendritic recordings were rejected if the series resistance exceeded 20 or 70 MΩ, respectively. Signals were lowpass filtered (DC to 10 kHz) and acquired by Axograph software at 50-100 kHz. At the termination of each recording, neuronal morphology and the placement of recording electrodes were recorded by fluorescence microscopy (QImaging). For nucleated patch recording, pipettes which had an open tip resistance of 2-4 MΩ when filled with the above internal solution were used to form high seal-resistance cell-attached recordings from visually identified pyramidal neurons, and the pipette capacitance was fully compensated. On successful excision of a nucleated patch, the patch was raised above the brain slice to the level of a prepositioned sylgard ball formed at the tip of a second pipette controlled by an independent manipulator. Current responses, recorded with a multi-clamp 700B amplifier (Molecular Devices), were evoked from a holding potential of −70 mV by the repeated delivery of voltage command steps (1000 repetitions, −5 mV, 7 ms). Recordings were lowpass filtered (DC to 30 kHz) and sampled at high frequency (250 kHz). High-resolution (60 x objective, 4 x optical magnification) photomicrographs were taken of each nucleated patch, allowing measurement of the major and minor axes and the calculation of surface area ^40^ At the termination of each recording the patch was ruptured, blown from the mouth of the pipette, and the recording pipette positioned to touch the sylgard ball where a high-resistance seal was formed. The voltage protocol was then repeated. The digital averages of current responses under these conditions were subtracted to remove any uncompensated residual pipette capacitance ^40^. Experimental data were analysed using Axograph, GraphPad Prism, SigmaPlot and Microscoft Excel. The voltage threshold of AP initiation was calculated as the point when the first derivative of somatically recorded APs exceeded 15 V/s. The recorded membrane potential values were adjusted for pipette offset, but not liquid junction potential.

### Microscopy and neuronal reconstruction

For accurate reconstruction, pyramidal neurons were filled with 0.5 % biocytin (wt / vol, 5 mg / ml, Sigma) and slices fixed in 4 % paraformaldehyde (pH 7.3 - 7.4) for 12 h. The slices were then reacted with 0.1 % streptavidin conjugated with Alexa Fluor-568 (Invitrogen) and 0.1 % Triton X-100 for 12 h at room temperature. Slices were subsequently reacted with DAPI (0.02 % in saline) for 30 min at room temperature, and then mounted and cover-slipped with Aqua Polymount (Polysciences) or Vectashield (Vector Laboratories). Optically sectioned (0.5 μm z step size) x-y photomicrograph montages of neuronal morphology were acquired using an inverted spinning-disk confocal microscope, either a W1 Yokogawa spinning disk module (3i), Zeiss Axio Observer Z1 microscope, Hamamatsu Flash4 sCMOS camera, and 40 x 1.2 NA C-Apochromat water immersion objective, or a Diskovery spinning disk module, Nikon TiE microscope, Zyla 4.2 sCMOS camera, and 40 x 1.15 NA Plan-Apochromat water immersion objective. Photomicrographs were acquired using 1 x 1 binning, with an exposure of 200-450 ms, set to allow the greatest dynamic range for all processes while not being so long as to over-bleach the fluorophore. Neurons were reconstructed manually and/or semi-automatically using a user-guided algorithm with Neurolucida 360 (MicroBrightField) at high digital magnification in 3 dimensions, and the reconstructions visualized and analysed with Neurolucida Explorer (MicroBrightField) software. Neurons possessing dendritic trees that appeared substantially truncated by the slicing procedure were excluded from further analysis. Dendritic diameter was measured using Neurolucida 360, where apical dendritic segments were semi-automatically traced with the user-guided rayburst crawl algorithm. For proximal apical dendrite segments, the segment began 15 μm from the center of the soma and continued for 20 μm. For distal apical segments (20 μm), the segment began 20 μm proximal to the main bifurcation of the apical dendrite (the nexus). Basal dendritic segments were manually traced to the first branch point or if a measured basal dendrite did not branch, it was measured for 20 μm or to termination, whichever was shorter. Traced segments were then loaded into Neurolucida Explorer and the average thickness of each segment calculated. For HL2/3 pyramidal neurons we also measured the average diameter of the distal 10 μm of proximal apical dendritic segments, to eliminate possible distortion of the diameter imposed by measurement of the soma-dendritic border. For illustrative purposes only, dendritic processes of reconstructions were thickened in Adobe Illustrator in order to provide clarity in text figures.

### Other statistics

No strategy for randomization and/or stratification was adopted. It was not possible to perform this study blinded. Inclusion and exclusion criteria for individual data types are detailed below.

Values reported are mean ± standard error of the mean (S.E.M.), unless otherwise stated, and n values represent the number of cells, unless otherwise stated. Statistical significance was defined when the probability of the data being observed under a null hypothesis was less than an alpha level of 0.05. Significance was tested with a two-tail Students’ t-test, with Welch’s correction being used for samples with unequal variance. The Mann Whitney test was used as a non-parametric test. Details of statistical tests can be found in the Results section. Statistical analysis was carried out using Prism 9 (Graphpad).

Only supragranular pyramidal neurons were included in the analysis. The laminar position of each neuron was determined by the merging of fluorescence images of biocytin-filled neurons with the nuclear counterstain DAPI, which was used to define neocortical lamination. Somatic or dendritic electrophysiological recordings were rejected if the somatic series resistance changed by >15 % during the experiment, or the dendritic series resistance changed by >40 %, or exceeded 20 or 70 MΩ, respectively. Recordings were also rejected if the observed recordings were highly unstable, or if neurons had a membrane potential that was depolarized relative to the norm.

## Acknowledgements

We are grateful to Rowan Tweedale for comments on a previous draft of the manuscript, and Rumelo Amor, Andrew Thompson and Arnaud Gaudin for their help with procedures for the imaging and reconstruction of neurons. We are grateful to Sue Engstrom and Karen Campbell (Brisbane Clinical Neuroscience Centre) for their help with project logistics. We are particularly grateful to the patients for their very generous donation. This work was supported by the National Health and Medical Research Council of Australia (GA59869 to SRW). Imaging was performed at the Queensland Brain Institute’s Advanced Microscopy Facility, generously supported by the Australian Government through the Australian Research Council LIEF grants LE100100074 and LE130100078.

## Author Information

### Affiliations

Queensland Brain Institute, The University of Queensland; Brisbane, Queensland, Australia

Helen M. Gooch, Tobias Bluett, Lisa Gillinder, Madhusoothanan B. Perumal, Hong D. Vo, Lee N. Fletcher, Stephen R. Williams

Mater Centre for Neurosciences, Mater Hospital; Brisbane, Queensland, Australia

Damian Amato, Robert Campbell, Lisa Gillinder, Jason McMillen, Jason Papacostas, Tony Tsahtsarlis, Martin Wood

Royal Brisbane and Women’s Hospital; Brisbane, Queensland, Australia

Michael J. Colditz, Rosalind L. Jeffree

### Contributions

Experimental: SRW, HMG, MBP and TB performed experiments. Analysis: SRW, HMG and MBP analysed electrophysiological data, HMG, TB, HDV, and LNF performed immunohistochemistry, confocal imaging, and neuronal reconstructions. Ethics and project management: LG and HMG. Surgery: JP, RLJ, MW, MJC, JM, TT, DA and RC provided human tissue. Conceptualization: SRW. Writing: SRW wrote the original draft manuscript, all authors participated in writing, reviewing, and editing of the manuscript.

Corresponding author. Stephen R. Williams

## Ethics declaration

### Competing interests

The authors declare no competing financial interests.

## Data and availability

All data are available in the main text and supplementary materials.

## Notes

### Competing Interest Statement

The authors have declared no competing interest.

### Summary of Updates

Typographical errors have been fixed

