## Supplementary Material for "High-fidelity dendritic sodium spike generation in human layer 2/3 neocortical pyramidal neurons"

Supplementary Figs. 1-8

Supplementary Tables 1-2

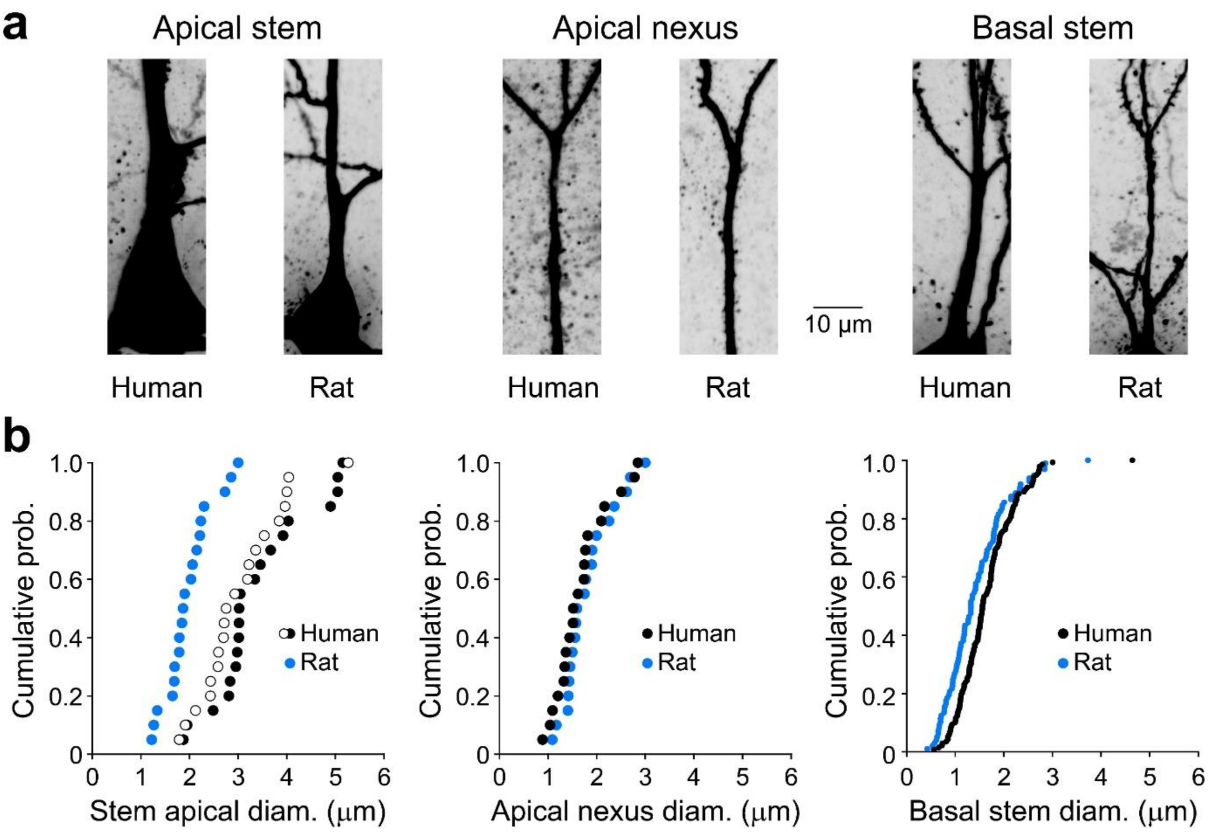


**Supplementary Fig. 1. Diameter of human and rat L2/3 pyramidal neuron dendrites**

**a,** Representative confocal images of dendritic segments in human and rat layer 2/3 pyramidal neurons. The indicated sites were selected as representative of average diameters. **b,** Pooled analysis of the diameter of apical and basal dendrites of human and rat layer 2/3 pyramidal neurons. The diameters of stem apical and basal dendrites were significantly different (apical stem: human= 3.43 ± 0.22 µm, rat= 1.98 ± 0.11 µm, Mann Whitney U= 28, *P*< 0.0001; basal stem: human= 1.66 ± 0.05 µm, rat= 1.44 ± 0.06 µm, Mann Whitney U= 5142, *P*= 0.0019). The open black symbols represent measurement of the diameter of stem apical dendrites in HL2/3 neurons over the distal 10 µm of the segment (apical stem: human (distal 10 µm) = 3.07 ± 0.19 µm, rat= 1.98 ± 0.11 µm, t-test, df= 38, *P*< 0.0001). Note, however, that the diameters of apical dendritic segments measured over a segment 20 µm proximal to the first main bifurcation of the apical dendritic tree (nexus) are similar between human and rat L2/3 pyramidal neurons (apical nexus: human= 1.69 ± 0.12 µm, nexus= 309 ± 54 µm from soma; rat= 1.82 ± 0.12 µm, nexus= 155 ± 20 µm from soma; t-test, df= 38, *P*= 0.4425).

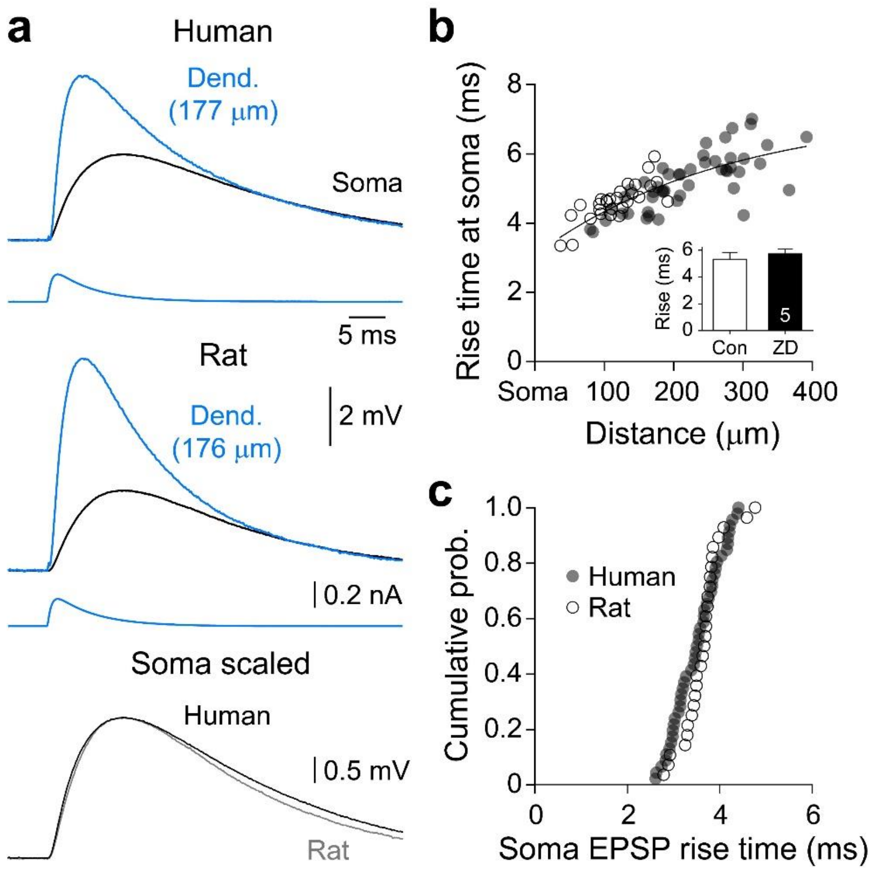


**Supplementary Fig. 2. Somatic rise time of dendritic simulated EPSPs**

**a,** Simulated dendritic EPSPs generate somatic voltage responses with similar rise times in human and rat L2/3 pyramidal neurons. Simultaneous recordings of somatic (black traces) and apical dendritic (blue traces) voltage are shown together with the driving current (lower traces). The lower overlain traces (soma scaled) show that the somatic rise-time of EPSPs generated at similar dendritic locations are nearly identical in a human and rat L2/3 pyramidal neuron. **b**, Quantification of the distance-dependent increase in the 10-90 % rise time of somatic sEPSPs generated at the indicated apical dendritic sites in HL2/3 (filled symbols) and RL2/3 (open symbols) pyramidal neurons. Data have been fit by an exponential function with a length constant of 279 μm. The inset shows that blocking HCN channels with ZD7288 (10 μM) in HL2/3 pyramidal neurons has little impact on the somatic rise time of dendritically generated sEPSPs. Bars represent mean ± S.E.M.. **c,** Cumulative probability distributions demonstrate the similar 10-90 % rise times of somatically generated and recorded sEPSPs in HL2/3 (filled symbols) and RL2/3 (open symbols) pyramidal neurons.

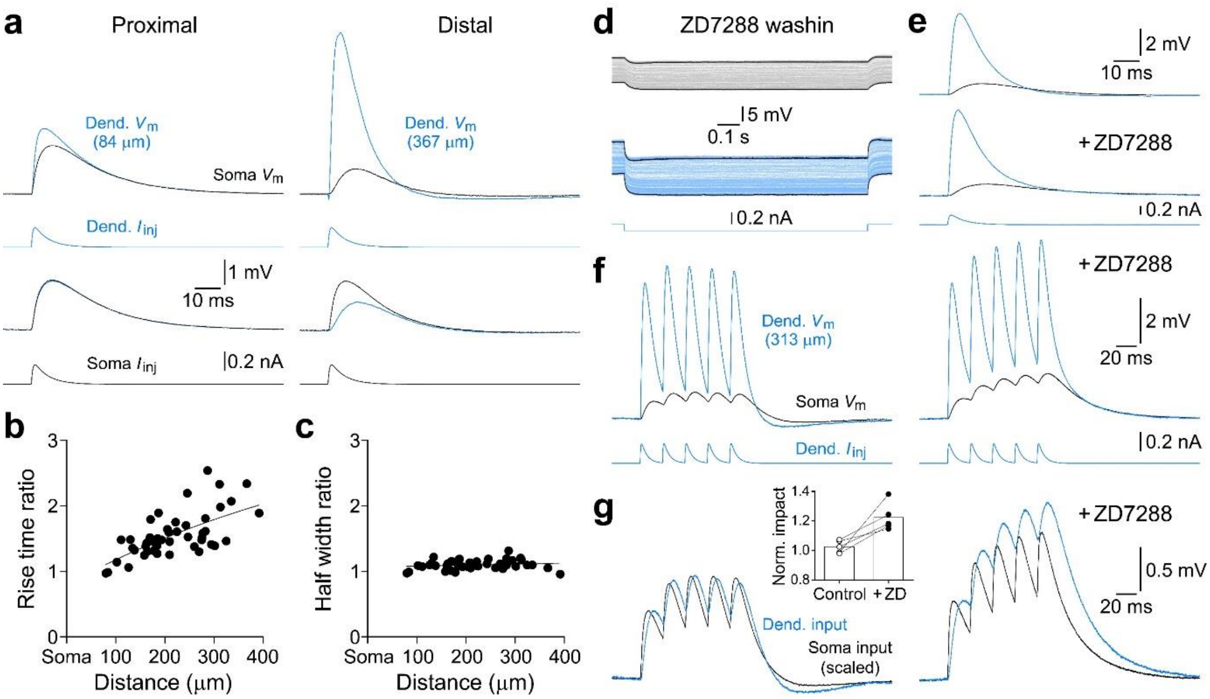


**Supplementary Fig. 3. HCN channels sculpt dendritic sEPSPs in human L2/3 pyramidal neurons**

**a,** Representative simultaneous somatic (black traces) and dendritic (blue traces) recordings of sEPSPs showing the distance-dependent pattern of dendro-somatic and somato-dendritic voltage attenuation. **b,c,** Quantification of the dendritic distance-dependent increase of the somatic rise time of sEPSPs, but the distance-dependent preservation of the somatic half-width of dendritically generated sEPSPs. Values represent the ratio of the kinetics of somatically recorded sEPSPs when generated at the soma and from the indicated dendritic sites in each tested HL2/3 pyramidal neuron. Data in panel b were fit with an exponential function with a length constant of 637 μm (line). Data in panel c were fit by linear regression, with a slope of 0.148 per mm (line). **d,** HCN channels powerfully control the resting membrane potential of HL2/3 pyramidal neurons. Simultaneous recording of somatic (grey traces) and dendritic (blue traces, 369 μm) voltage responses generated by a negative dendritic current step (lower trace) under control and during the application of ZD7288 (10 μM). The overlain black traces represent digital averages compiled under control conditions and in the presence of ZD7288. Pooled results demonstrated that the application of ZD7288 significantly hyperpolarized the resting membrane potential of HL2/3 neurons (control soma= -72.6 ± 0.8 mV, ZD soma= -80.5 ± 1.8 mV, n= 5; paired t-test, P= 0.0052; control dendrite= -70.0 ± 1.5 mV, ZD dendrite= -77.7 ± 2.7 mV; paired t-test, P< 0.0151; average distance from the soma: 255 ± 39 μm). **e,** The pharmacological blockade of HCN channels eliminated the cross-over in the decay phase of dendritic (blue traces) and somatic (black traces) sEPSPs delivered 312 μm from the soma (lower trace). **f,** Temporal summation of sEPSP trains simultaneously recorded at the dendritic site of generation (blue traces) and the soma (black traces). Note the greater temporal summation of sEPSP trains when HCN channels were blocked with ZD7288 (10 μM). **g,** The somatic impact of dendritic sEPSP trains is controlled by HCN channels. The amplitude of the first somatically recorded sEPSP of the train has been peak scaled; same neuron as in panels e and f. The inset shows quantification of the enhanced summation of dendritically generated sEPSPs in ZD7288.

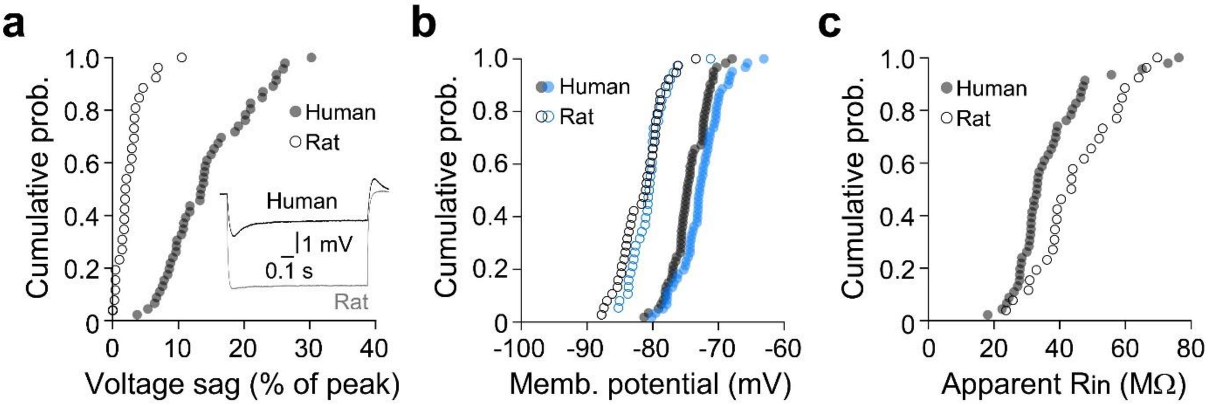


**Supplementary Fig. 4. Electrophysiological properties of human and rat L2/3 pyramidal neurons**

**a,** Somatic voltage sag (% of peak) apparent during negative voltage responses (inset) is greater in HL2/3 pyramidal neurons (Mann Whitney U= 23, *P*< 0.0001). **b,** Distribution of somatic (black) and apical dendritic (blue) membrane potential of human (filled symbols) and rat L2/3 (open symbols) pyramidal neurons; distributions are significantly different between species (soma: t-test, df= 97, *P*< 0.0001; dendrite: *P*< 0.0001). **c,** The peak somatic apparent input resistances of human and rat L2/3 pyramidal neurons are distinct (Mann Whitney U= 344, *P*< 0.0026).

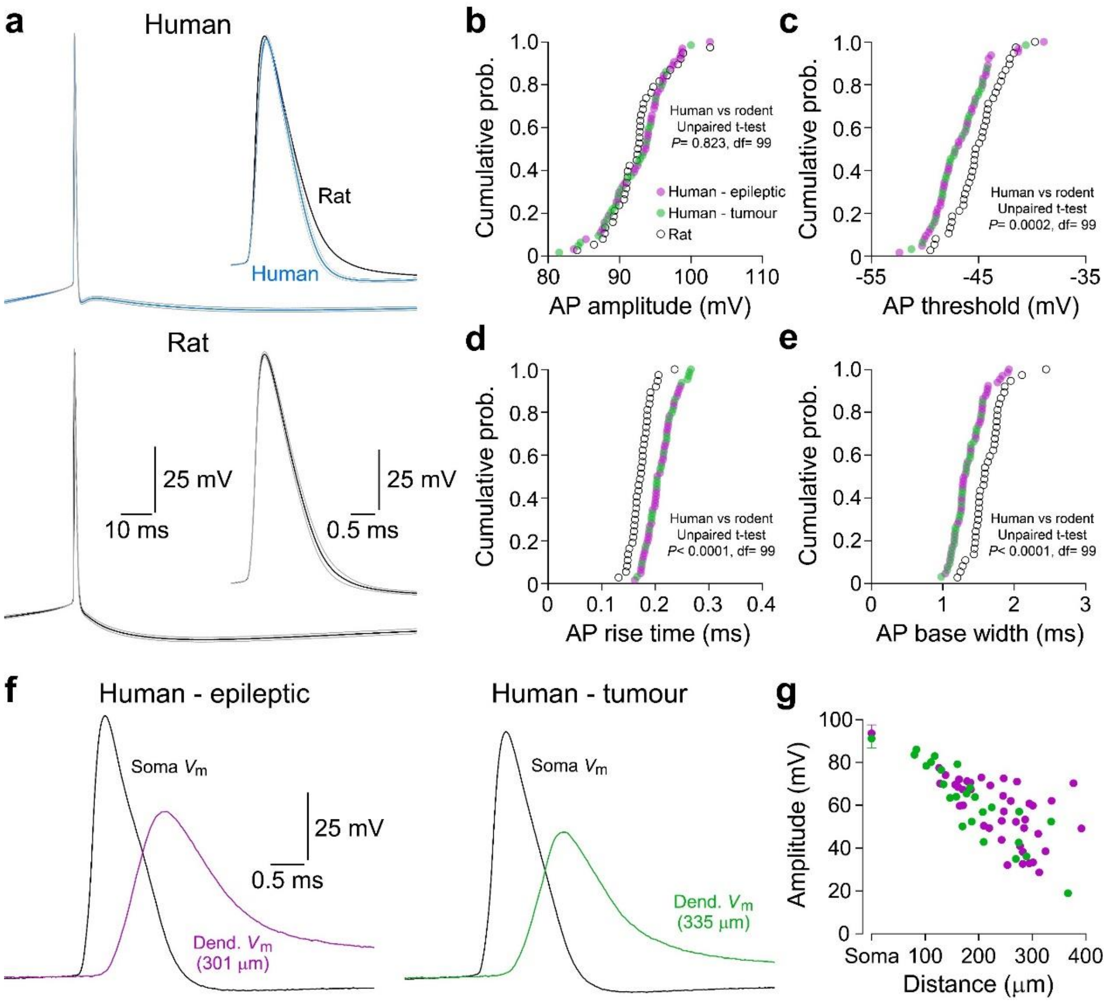


**Supplementary Fig. 5. Properties of APs and BPAPs in human and rat L2/3 pyramidal neurons**

**a,** Averaged waveform of somatically recorded APs from 51 HL2/3 (blue) and 35 rat (black) L2/3 pyramidal neurons. APs were generated by the presentation of somatic rheobase positive current steps. The grey lines represent error around the mean (± 2 S.E.M.). Note the characteristic differences in AP after-hyperpolarizing potentials. The averaged APs are shown inset at a faster time-base to illustrate the rise- and decay time course. **b-e,** Cumulative probability distributions of the indicated properties of APs recorded from human and rat L2/3 pyramidal neurons. Human neurons are delineated by etiology. Note that the properties of APs recorded from patients who underwent surgery for the removal of sub-cortical brain tumours (green symbols) or the alleviation of refractory epilepsy (magenta symbols) intermingle throughout the distributions. For each panel the results of statistical comparison between human and rat L2/3 pyramidal neurons are shown. **f,** Representative waveforms of somatic APs and simultaneously recorded dendritic BPAPs in HL2/3 pyramidal neurons recorded from patients who had undergone surgery for the removal of a sub-cortical brain tumour or the alleviation of refractory epilepsy. **g,** Distance-dependency of the amplitude of BPAPs in HL2/3 pyramidal neurons delineated by patient etiology. The average amplitude of somatically recorded APs (± S.D.) is shown.

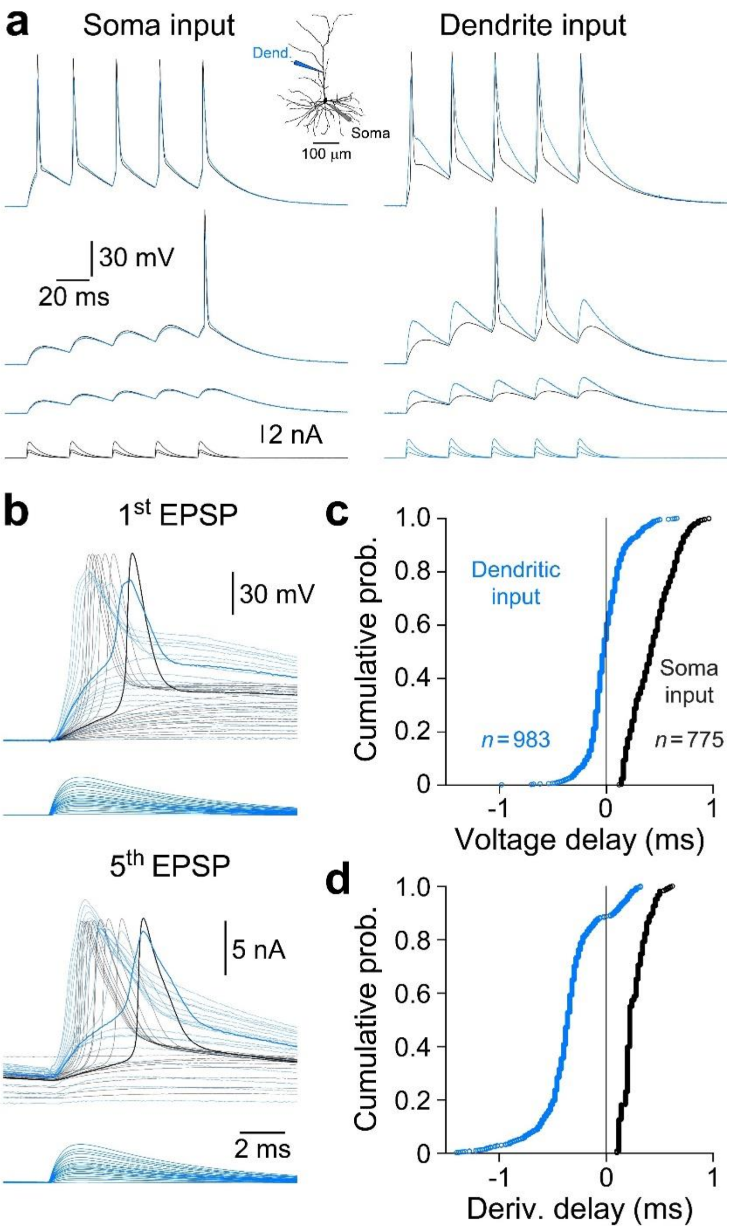


**Supplementary Fig. 6. Dendritic spikes are generated at high frequencies in rat L2/3 pyramidal neurons**

**a,** Repetitive dendritic spike generation is driven by simulated excitatory postsynaptic current trains (lower overlain traces) in a RL2/3 pyramidal neuron. The morphology of the neuron and placement of recording electrodes are shown inset. **b,** Baselined sEPSPs generated by the first and fifth sEPSP of the train at a faster time base (same neuron as a). Note that dendritic spikes drive AP firing over a wide suprathreshold range. **c,d,** Cumulative probability distributions of the timing of all regenerative events evoked by dendritic (blue) and somatic (black) sEPSP trains in RL2/3 pyramidal neurons. The time difference was calculated at peak voltage of events in c, and by the time difference between the peak amplitude of the first derivative of events in d. Note the clear disparity between timing relationship, and the preponderance of APs driven by dendritic spikes (negative times) when sEPSP trains were presented at dendritic sites.

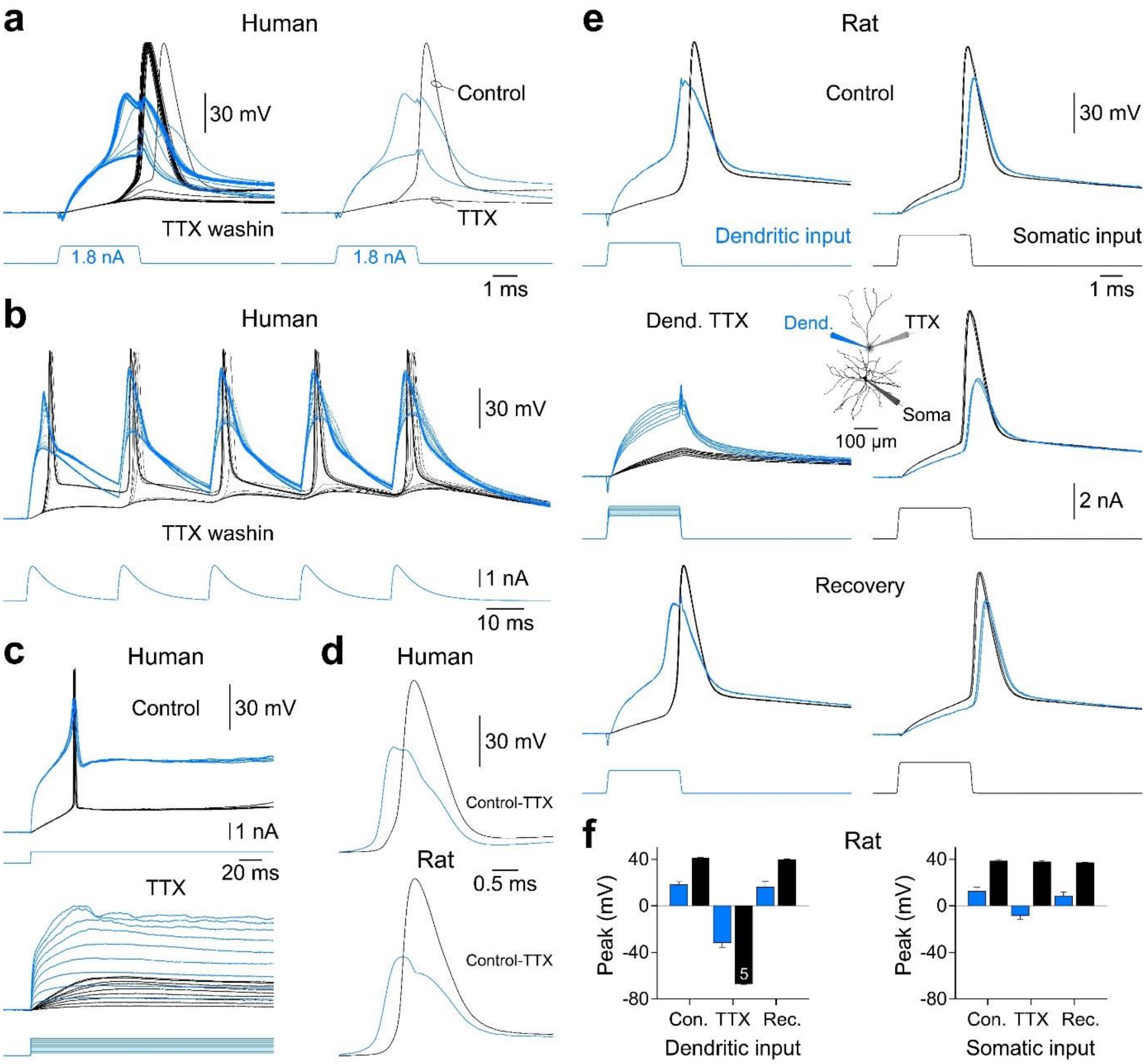


**Supplementary Fig. 7. Dendritic sodium spikes in rat and human L2/3 pyramidal neurons**

**a,** The bath application of TTX (1 μM) blocks regenerative dendritic spikes evoked by a threshold short dendritic positive current pulse in a HL2/3 pyramidal neuron. Note the blockade of dendritic spikes and APs as TTX is washed into the bath. The traces on the right side show the waveform of a dendritic spike and coupled AP under control and the electrotonic responses generated in the presence of TTX. **b,** Dendritic spikes generated repetitively by a train of simulated dendritic excitatory postsynaptic currents are blocked by the bath application of TTX in a HL2/3 pyramidal neuron. Note the abolition of dendritic spikes and APs as TTX is washed into the bath. **c,** Overlain traces of dendritic spikes generated in response to threshold long dendritic current steps (upper traces) are blocked by the bath application of TTX (1 μM, lower overlain traces). In the presence of TTX regenerative activity is abolished across a wide positive voltage range. **d,** Representative waveforms of TTX subtracted dendritic spikes and APs evoked by threshold positive current steps in a human and rat L2/3 pyramidal neuron. **e,** Overlain voltage traces show dendritic (blue) and somatic (black) responses evoked by brief dendritic or somatic positive current steps (lower traces) in a RL2/3 pyramidal neuron. The transient local dendritic application of TTX (2 μM) blocked the initiation of dendritic spikes but did not alter the characteristics of APs generated by somatic positive current pulses. A reconstruction of the morphology of the RL2/3 pyramidal neuron is shown inset, illustrating recording and TTX application sites. **f,** Pooled data describing the blockade of dendritic spikes and the reduction of the amplitude of BPAPs by the local dendritic application of TTX. Bars represent mean ± S.E.M. and n refers to the number of cells.

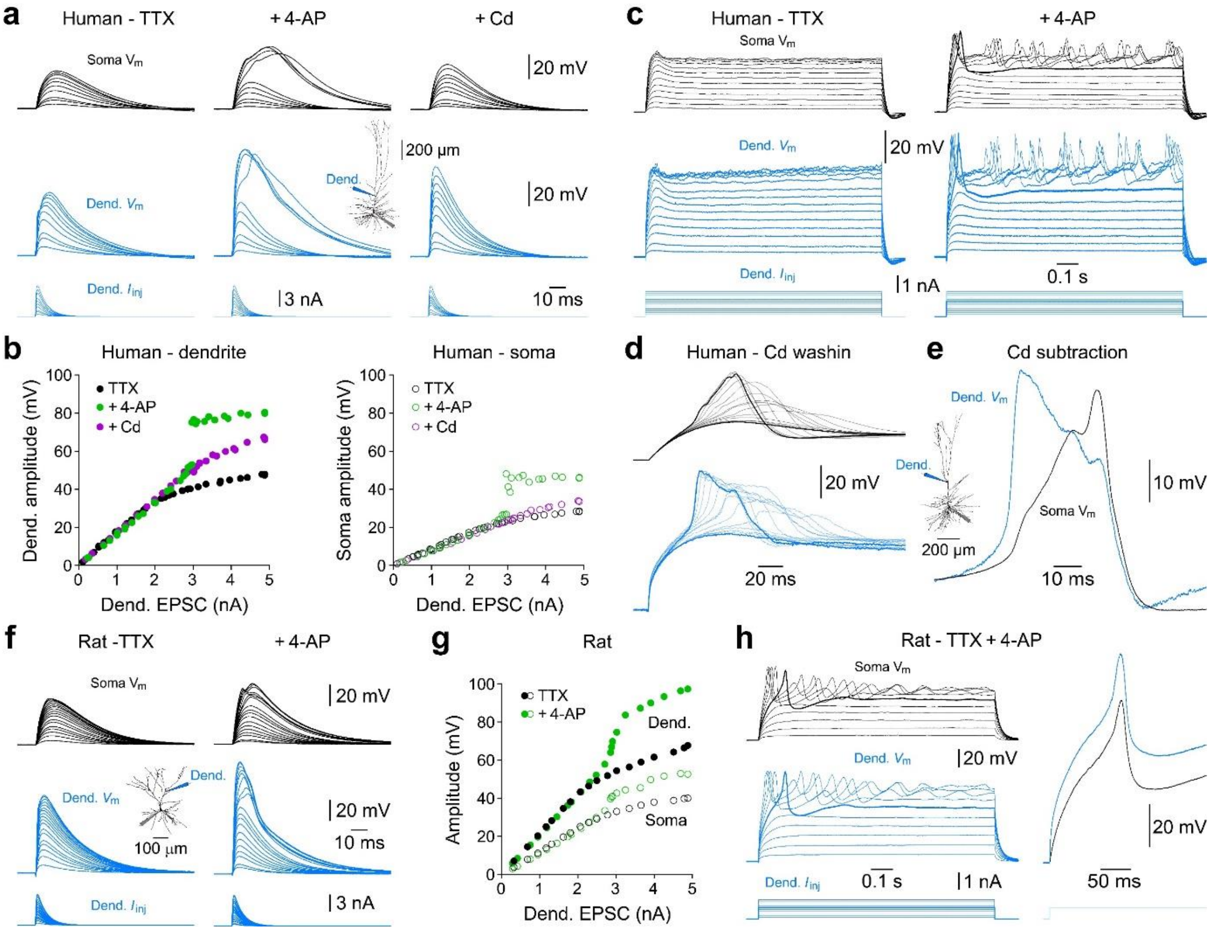


**Supplementary Fig. 8. Pharmacological blockade of potassium channels unmasks dendritic calcium electrogenesis in human and rat L2/3 pyramidal neurons**

**a,** Simultaneous somatic and apical dendritic recording of families of simulated dendritic EPSPs recorded under the indicated conditions from a HL2/3 pyramidal neuron (morphology and recording sites inset). Note the absence of dendritic regenerative activity when sodium channels were blocked with tetrodotoxin (TTX, 1 μM), but the generation of powerful dendritic electrogenesis following the addition of the voltage-gated potassium channel antagonist 4-amino pyridine (4-AP, 5 mM). Under these conditions dendritic regenerative activity was blocked by the addition of the broad-spectrum voltage-gated calcium channel antagonist cadmium (Cd, 100 μM). **b**, Measurement of the peak amplitudes of dendritic sEPSPs at dendritic site of generation (left graph) and the soma (right graph) under the indicated conditions, same HL2/3 pyramidal neuron as panel a. **c**, Families of voltage responses evoked by the injection of long steps of positive current at the dendritic recording site (160 μm from the soma) of a HL2/3 pyramidal neuron under the indicated conditions. Note the absence of dendritic regenerative activity in the presence of TTX (1 μM), but the unmasking of repetitive regenerative dendritic activity by the bath addition of 4-AP (5 mM). The threshold response is demarked by bold traces. **d**, The bath application of cadmium (100 μM) blocks dendritic electrogenesis recorded in the presence of TTX (1 μM) and 4-AP (5 mM) from a HL2/3 pyramidal neuron (morphology and recording sites inset). The bold traces delineate control (TTX and 4-AP) and cadmium conditions. **e**, The waveform of cadmium subtracted dendritic calcium electrogenesis recorded at dendritic site of generation and the soma, same HL2/3 pyramidal neuron as shown in panel d. **f**, Simultaneous somatic and apical dendritic recording of families of simulated dendritic EPSPs recorded under the indicated conditions from a RL2/3 pyramidal neuron (morphology and recording sites inset). Note, the absence of dendritic regenerative activity when sodium channels were blocked with tetrodotoxin (TTX, 1 μM), but the generation of dendritic electrogenesis following the blockade of voltage-gated potassium channels with 4-amino pyridine (4-AP, 5 mM). **g**, Measurement of the peak amplitudes of sEPSPs at dendritic site of generation (filled symbols) and the soma (open symbols) under the indicated conditions. **h**, Families of voltage responses evoked by the injection of long steps of positive current at the dendritic recording site of the same RL2/3 pyramidal neuron in the presence of TTX (1 μM) and 4-AP (5 mM). Note the generation of robust calcium electrogenesis. The threshold response is delineated by the bold traces and shown at a faster time-base inset.

### Supplementary Table 1. Information on patients with refractory epilepsy

| # | Age | Sex | Diagnosis | Age of epilepsy onset | Recent antiepileptic drugs | Anesthetic | Paralytic agent | Resected brain region |
|---|---|---|---|---|---|---|---|---|
| 1 | 55 | M | Epilepsy - hippocampal sclerosis | 20 | LTG, BRV, CBZ, CLB | PRO, REM, SEV, FEN | ROC | L-temporal |
| 2 | 25 | F | Epilepsy - mesial temporal sclerosis | 5 | TOP, LTG, LCM | PRO, REM | ROC | L-temporal |
| 3 | 22 | M | Epilepsy | 15 | OXC, BRV, LCM, ZNS, CLB | PRO, REM, FEN, MID | ROC | L- temporal |
| 4 | 32 | M | Epilepsy - focal cortical dysplasia | childhood | LEV, LCM, LTG | PRO, REM, FEN | VEC | R-temporal |
| 5 | 42 | M | Epilepsy - mesial temporal sclerosis | 24 | PER, CBZ, LCM, BRV | PRO, REM | ROC | R-temporal |
| 6 | 17 | M | Epilepsy - dysembryoplastic neuroepithelial tumor (DNET) | 15 | PER, OXC, BRV, LTG, CLB | PRO, REM, FEN | VEC | L-temporal |
| 7 | 54 | M | Epilepsy - mesial temporal sclerosis | 39 | LCM, LEV | PRO, REM | ROC | L-temporal |
| 8 | 23 | M | Epilepsy - hippocampal sclerosis | 13 | CLB, CBZ | PRO, REM | ROC | R-temporal |
| 9 | 22 | F | Epilepsy - focal cortical dysplasia-like | 5 | ZNS, BRV, LCM | PRO, REM, FEN | VEC | L-temporal |

Abbreviations

Antiepileptic agents: LEV – levetiracetam, LTG – lamotrigine, CBZ – carbamazepine, LCM – lacosamide, OXC – oxcarbazepine, CLB – clobazam, ZNS – zonisamide, BRV – brivaracetam, PER – perampanel, TOP – topiramate.

General anesthesia maintenance: PRO – propofol, REM – remifentanil, SEV – sevoflurane, FEN – fentanyl, MID – midazolam, LIG – lignocaine.

Paralytic agents: ROC – rocuronium, VEC – vecuronium.

L – left, R – right.

### Supplementary Table 2. Information on patients with subcortical tumours

| # | Age | Sex | Diagnosis | Recent antiepileptic drugs | Anesthetics | Paralytic agent | Resected brain region | 5-ALA |
|---|---|---|---|---|---|---|---|---|
| 1 | 70 | F | Tumour - glioblastoma | LEV (during surgery) | PRO, REM, FEN, MID | VEC | L-temporal | No |
| 2 | 50 | F | Tumour - glioblastoma | LEV | PRO, REM | ROC | R-temporal | Yes |
| 3 | 69 | F | Tumour - glioblastoma | LEV | PRO, REM | ROC | L-temporal | Yes |
| 4 | 37 | M | Focal cortical dysplasia | LTG, LEV, PER | PRO, REM, LIG | ROC | L-temporal | No |
| 5 | 59 | M | Tumour - glioma | N/A | PRO, REM, FEN, MID | VEC | R-temporal | No |
| 6 | 60 | F | Tumour - meningioma | N/A | PRO, REM, LIG | VEC | R-temporal | No |

Abbreviations

Antiepileptic agents: LEV – levetiracetam, LTG – lamotrigine, PER – perampanel.

General anesthesia maintenance: PRO – propofol, REM – remifentanil, FEN – fentanyl, MID – midazolam, LIG – lignocaine.

Paralytic agents: ROC – rocuronium, VEC – vecuronium.

Imaging agent: 5-ALA – 5-aminolevulinic acid.

L – left, R – right.
